# Genomic acquisitions in emerging populations of *Xanthomonas vasicola* pv. *vasculorum* infecting corn in the U.S. and Argentina

**DOI:** 10.1101/587915

**Authors:** Alvaro L Perez-Quintero, Mary Ortiz-Castro, Guangxi Wu, Jillian M. Lang, Sanzhen Liu, Toni A Chapman, Christine Chang, Janet Ziegle, Zhao Peng, Frank F. White, Maria Cristina Plazas, Jan E. Leach, Kirk Broders

## Abstract

*Xanthomonas vasicola* pv. *vasculorum (Xvv)* is an emerging bacterial plant pathogen that causes bacterial leaf streak on corn. First described in South Africa in 1949, reports of this bacteria have greatly increased in the past years in South America and in the U.S., where it is now present in most of the corn producing states. Phenotypic characterization showed that the emerging U.S. and South American X*vv* populations may have increased virulence in corn compared to older strains. To understand the genetic mechanisms behind the increased virulence in this group, we used comparative genomics to identify gene acquisitions in *Xvv* genomes from the U.S. and Argentina. We sequenced 41 genomes of *Xvv* and the related sorghum-infecting *X. vasicola* pv. *holcicola* (*Xvh).* A comparison of all available *X. vasicola* genomes showed the phylogenetic relationships in the group and identified clusters of genes associated with the emerging *Xvv* populations. The newly acquired gene clusters showed evidence of horizontal transfer to *Xvv* and included candidate virulence factors. One cluster, in particular, corresponded to a prophage transferred from *Xvh* to all *Xvv* from Argentina and the U.S. The prophage contains putative secreted proteins, which represent candidates for virulence determinants in these populations and await further molecular characterization.

## Introduction

In the U.S., bacterial leaf streak of corn (BLS) was first observed in Nebraska in 2014 and became widespread by 2016 (Korus et al. 2017). The disease is present in dent corn and popcorn producing regions of Colorado, Kansas and Nebraska, with several fields reporting disease incidence levels above 90% and disease severity reaching greater than 50% of leaf area infected (Broders 2017). The disease has now been found in most of the corn growing region of the U.S. including Illinois, Iowa and Nebraska, which are the top three corn producing states in the U.S. (Korus et al. 2017; USDA-NASS 2017). Given the large number of acres and economic importance of corn production in the U.S., there are important implications to the emergence and spread of this new disease. Thus, understanding how this disease originated and what favors its spread is crucial to prevent future losses.

Caused by *X. vasicola* pv. *vasculorum (Xvv*), BLS was first described in 1949 on corn in South Africa (Dyer 1949), but prior to 2016 it had not been documented in any other country. It is unknown how this organism was introduced to the U.S. or if it was already present in a latent state. The only other report of BLS of corn outside of South Africa and the U.S. was in Argentina and Brazil in 2017 and 2018 (Plazas et al. 2017, Leite et al. 2019). While the official report of the disease in Argentina is relatively recent, the symptoms of BLS were first observed in 2010 in the Cordoba province and have since spread to nine other corn-producing provinces, including provinces that border the corn growing regions in Brazil and Paraguay (Leite et al. 2019; Plazas et al. 2017). It is still unclear why *Xvv* continues to spread in the Americas or how severe future epidemics may become. However, it does appear that *Xvv* may have been present in Argentina prior to reports of this pathogen in the U.S.

A significant amount of confusion existed around the taxonomic classification of this bacterium. The nomenclature had gone through several changes, from *X. campestris* pv. *zeae* to *X. vasicola* pv. *zeae* to its current designation as *X. vasicola* pv. *vasculorum (*Lang et al. 2017; Coutinho and Wallis 1991; Sanko et al. 2018; Bradbury 1986; Qhobela, Claflin, and Nowell 1990). The *X. vasicola* species is now divided into five groups including three named pathovars: 1) *Xvv* infecting corn and sugarcane, 2) *X. vasicola* pv. *holcicola (Xvh)*, commonly infecting sorghum, 3) *X. vasicola* pv. *musacearum (Xvm)* infecting enset and banana, 4) a group of strains isolated from *Tripsacum laxum*, and, 5) strains isolated from areca nut (previously *X. campestris* pv. *arecae*) (Lang et al. 2017) (Studholme et al., this issue).

The term pathovar refers to a strain or set of strains with the same or similar characteristics, differentiated at the infrasubspecific level from other strains of the same species or subspecies on the basis of distinctive pathogenicity to one or more plant hosts (Young et al. 1991). While the named pathovars of *X. vasicola* seem to have defined host preferences, their host ranges may be broader than initially claimed. *Xvh* and *Xvv*, in particular, may have overlapping host ranges. Under laboratory conditions, isolates of *Xvv* from corn and sugarcane caused disease on corn, sugarcane and sorghum, but were most virulent on corn and sugarcane (Lang et al. 2017). Similarly, when infiltrated into leaves, *Xvh* infected corn, sorghum and sugarcane, but caused more disease on sorghum (Lang et al. 2017). *Xvv* has not been isolated from sorghum, while *Xvh* has occasionally been isolated from corn in the field (Moffett 1983; Péros et al. 1994). Upon inoculation in the greenhouse, *Xvv* isolates from the U.S. can infect 16 hosts, mostly monocots such as rice, oats and big blue stem and the dicot yellow nutsedge (Hartman et al. this issue). Field studies confirmed these results for big blue stem and bristly foxtail as hosts in a natural inoculum system (Hartman et al. this issue). At least two host jumps have been hypothesized for *X. vasicola*, i.e. from grasses to banana (Tushemereirwe et al. 2004) and from sugarcane to *Eucalyptus* spp, a dicot (Coutinho et al. 2015), suggesting a remarkable adaptive ability for the species.

The appearance and spread of *Xvv* in the U.S. and Argentina was rapid. How and why the populations expanded so quickly in two countries located on the opposite side of the equator at approximately the same time, while the disease remained rarely documented in South Africa during the same time period, is intriguing. Reasons for these disparate observations could include the occurrence of more favorable environmental conditions and susceptible corn germplasm in the Americas versus South Africa, and/or, as we hypothesize here, the acquisition of genetic features that favored infection or spread or virulence. In this study, we employed a comparative genomics approach to identify genetic changes associated to these emerging populations.

## Results

### U.S. *Xvv* strains are closely related to each other and to *Xvv* strains from Argentina

Draft genome assemblies were generated for fifteen strains collected in 2016 in the states of Colorado, Iowa, Kansas and Nebraska. Draft genome sequences were also prepared for *Xvv* isolates from South Africa (2 strains from corn) and Argentina (7 strains), and for *Xvh* isolates from the U.S. (1 strain, from sorghum) and Australia (8 strains from sorghum, 3 from corn). Additionally, fully assembled genomes were derived for three U.S. *Xvv* isolates isolated from corn, one sugarcane *Xvv* isolate from Zimbabwe, and one sorghum *Xvh* isolate from Mexico, totaling 41 new genomes (Supplementary Table 1).

A phylogenetic maximum-likelihood tree was determined on the basis of the pan-genome SNPs (Gardner, Slezak, and Hall 2015), using the newly sequenced genomes as well as all available *Xanthomonas vasicola* genomes, adding *Xanthomonas oryzae* pv. *oryzae* PXO99A as an outgroup (94 genomes in total) (Supplementary Table 1). The tree reveals division of the four main groups/pathovars in the species: *holcicola (Xvh), musacearum (Xvm), vasculorum (Xvv)* and an unnamed group of strains collected from *Tripsacum laxum* (here referred to as simply *Xv)* (Figure 1 A). Most *Xvv* strains from corn formed a closely-related group, separated from strains isolated from sugarcane, with the exception of strain NCPPB206, a weakly virulent isolate from South Africa collected in 1948 (Lang et al. 2017; Dyer 1949). *Xvv* strains from the U.S. tended to group together (one exception NE-7 from Nebraska), and seemed to be more closely related to strains from Argentina than to strains from South Africa (Figure 1 A).

**Figure 1.**
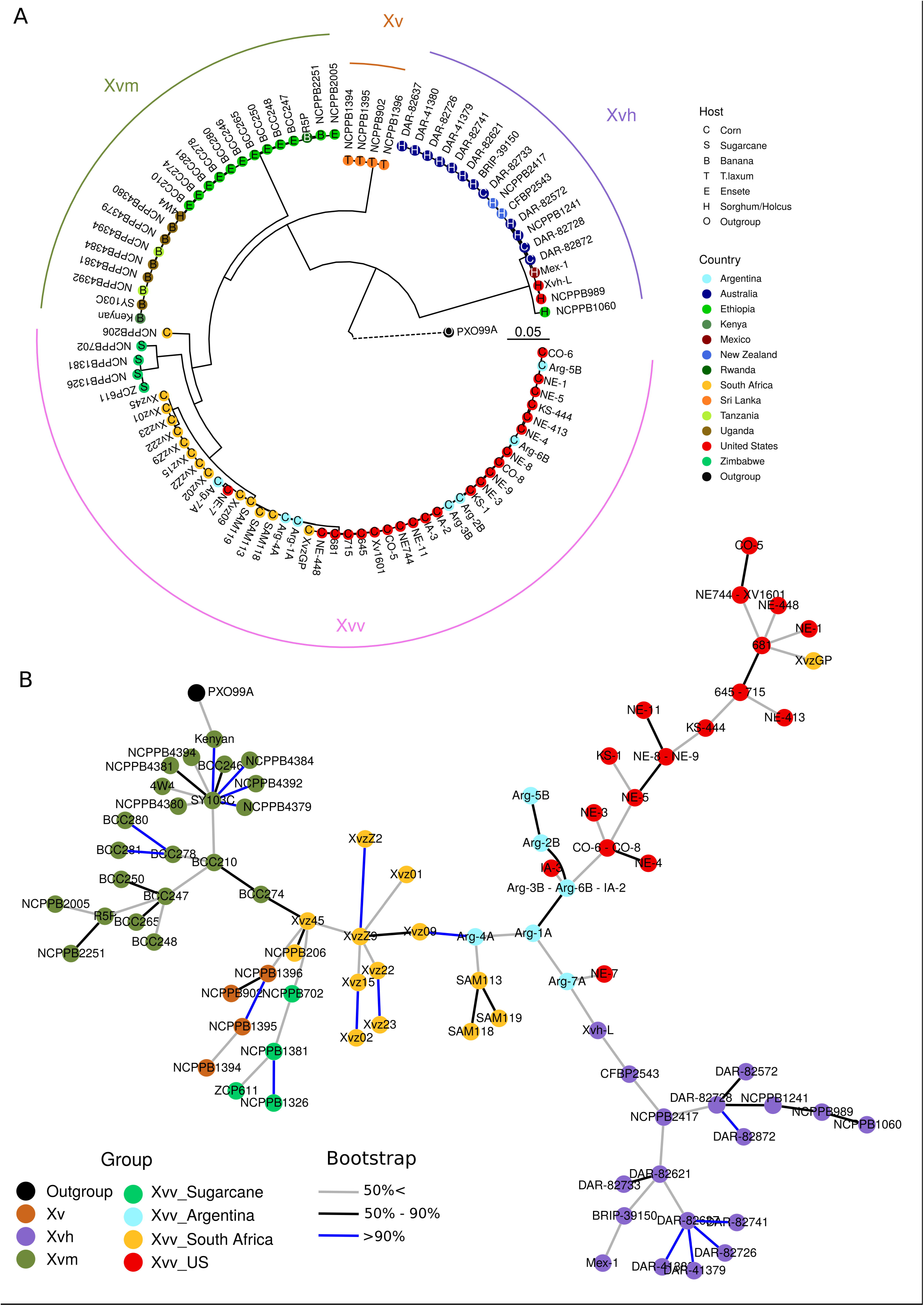
Phylogeny of *X. vasicola* strains. A) Parsimony tree based on pan-genome SNPs from 91 draft and fully sequenced *X. vasicola* genomes built using kSNP3 (Gardner, Slezak, and Hall 2015). Four main pathovars/groups are indicated by solid colored lines. Colors in tree tips indicate country of isolation and tip letters indicate the plant host of the isolate. Bar shows the tree scale. Dotted line to the out-group (*X. oryzae* pv. *oryzae*) indicates this distance was scaled down tenfold to improve visualization. Colors in the tip letters (black or white) are for readability and do not indicate any feature. 74% of the nodes had support over 70% (as calculated by kSNP3 (Gardner, Slezak, and Hall 2015). B) Consensus minimum spanning tree based on core genome SNPs with PXO99A as a reference. Circle colors indicate seven groups of interest of *X. vasicola.* The consensus tree was bootstrapped 500 times and edge colors indicate bootstrap percentages.

Similar groupings were seen in phylogenetic trees based on core-genome SNPs (Supplementary Figure 1 A), multi-locus sequence alignments (MLSA) from house-keeping genes or core-genome genes (Supplementary Figure 1), and average nucleotide identity (Supplementary Figure 2). Additionally, minimum spanning trees (MSTs) based on core-genome SNPs showed the same main groups but also revealed possible relations between the different corn *Xvv* groups. Unlike phylogenetic trees, MSTs only assume identity based only upon state (SNP) similarity and not upon relationships to hypothetical ancestors (Salipante and Hall 2011). By MST, Argentinian *Xvv* strains form a central cluster connected to U.S. and South African *Xvv* strains but also to *Xvh*, which indicates a possible transmission path whereby the Argentinian *Xvv* population is a link between the U.S. and South African ones, and possibly also received genetic material from *Xvh* (Figure 1B).

**Figure 2.**
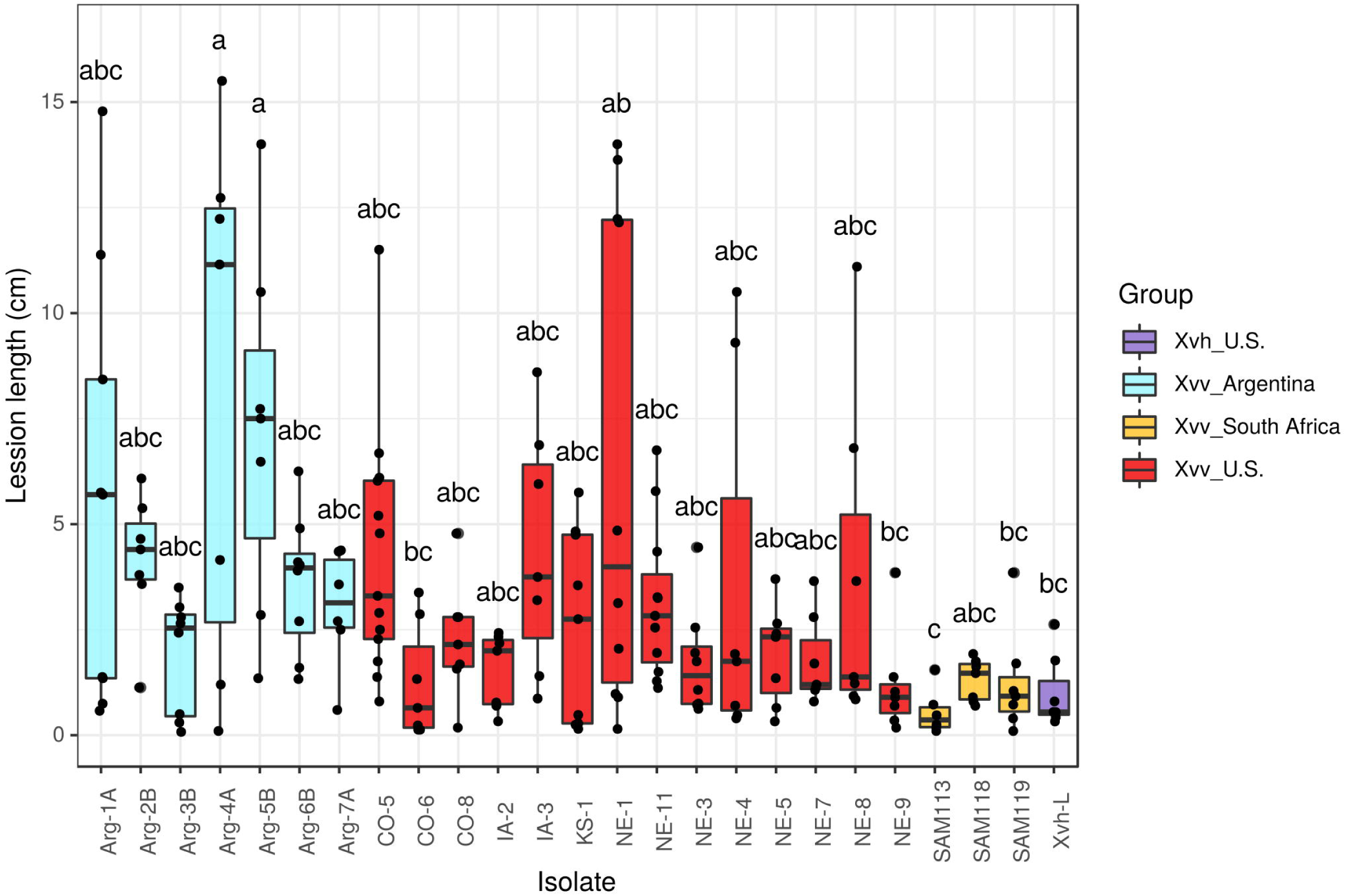
Phenotypic characterization of *Xvv* strains. Disease caused by *Xvv* and *Xvh* on corn (hybrid P1151). Three week-old plants were infiltrated with 10^8^ CFU mL^−1^ of each isolate, and disease was assessed at seven days post inoculation (dpi). Lesion lengths indicate expansion beyond the infiltration site. The experiment was replicated three times and combined data from all replications is shown here. Letters designate significance groups at *p* value <0.0001 using one-way ANOVA+Tukey’s HSD using square root transformation of data and sample size of at least seven replicates per isolate (90% statistical power).

Disease phenotyping showed that contemporary *Xvv* isolates from the US and Argentina tend to cause more severe symptoms on corn (hybrid P1151) than older *Xvv* isolates form South Africa and contemporary *Xvh*, although with high variation (Figure 2). Suggesting the recent epidemic may be associated to a gain of virulence in the American populations. These results however may be dependent on the host genotype since more severe symptoms have been reported for *Xvh* and South African *Xvv* in another corn variety (Lang et al. 2017).

### U.S. *Xvv* genomes contain clusters of genes often absent in other *X. vasicola* genomes

To identify genes associated to the emerging U.S. *Xvv* population, we identified ortholog groups among all proteins. Overall, 3616 genes were present in all the *X. vasicola* groups (Supplementary Figure 3), and the core genome (orthologs present in all strains) was 2755 genes (Supplementary Figure 3). Orthology distribution analysis in the different groups found that 44 genes were exclusive to corn *Xvv*. No genes were found to be exclusive to *Xvv* from the U.S., but 19 genes were shared exclusively with Argentinian strains. Sixty-three genes were shared between *Xvh* and Argentinian and U.S. *Xvv* strains (Supplementary Figure 3).

**Figure 3.**
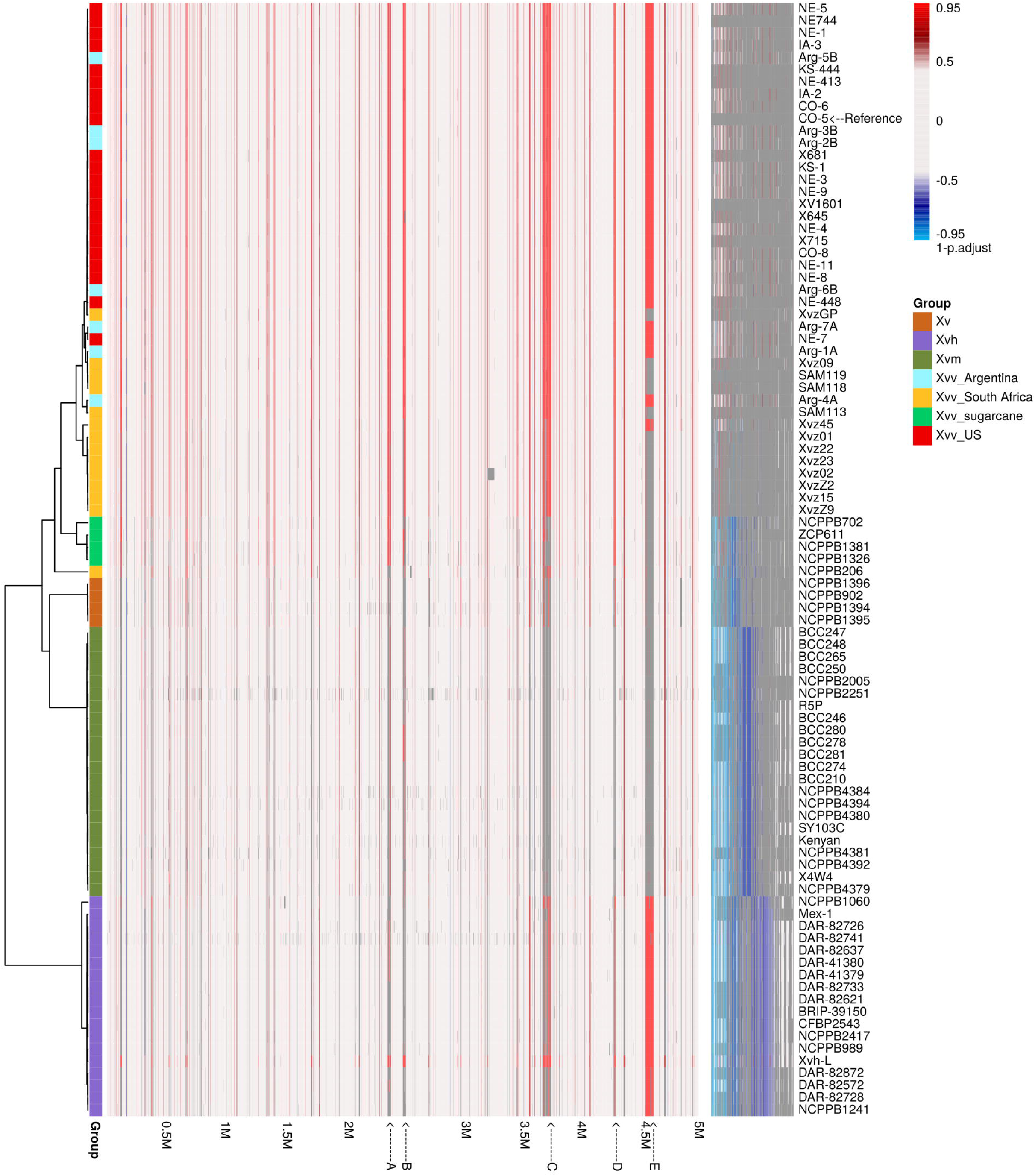
Over and under-represented genes in U.S. *Xvv.* Heatmap shows presence (white or colored) or absence (grey) of all ortholog groups found in at least 10 strains in the genome set, each vertical line represents an ortholog group (4912 total). A hypergeometric test was applied to each group to determine over or under representation in the group of U.S. *Xvv* strains versus all other groups. Colors in the heatmap (blue to red) indicate 1 - the adjusted *p* value for these tests. Negative values indicate the result of an under-representation test (adjusted *p* value * −1). The vertical lines in the heatmap are ordered according to the presence of each ortholog group in the genome of the U.S. *Xvv* strain CO-5, numbers below the heatmap (1M to 5M) indicate the position (in million base pairs) of the genes in the reference genome. Vertical lines after the 5M mark are orthologs not present in CO-5 ordered according to their frequency in other genomes. Bars at the left of the heatmap indicate the group of the strain as in Figure 1B. The dendrogram to the left corresponds to a MLSA tree based on all orthologs (Supplementary Figure 1). Five gene clusters of over-represented genes (A-E) were identified and indicated with letters below the heatmap, these clusters defined as groups of at least 10 genes found less than 5 Kb from each other in the CO-5 genome and with an adjusted *p* value for over-representation <0.05 (0.95 in the figure).

To uncover more genes associated with (but not necessarily exclusive to) the U.S. *Xvv* population, we performed a hyper-geometric test for each ortholog group. In this test we compared the presence and copy number of each gene in the U.S. *Xvv* genomes against 100 sets of randomly selected genomes from the other groups (Supplementary Figure 4). The test identified 278 genes that were over-represented in U.S. *Xvv* and 44 genes that were underrepresented (Figure 3, Supplementary Table 2). The Xvv U.S.-associated genes in the reference genome Xvv strain CO-5 were often grouped together, indicating sub-genomic regions are associated with the population. Five clusters of over-represented genes (named A to E) were identified (Figure 3). The clusters contain 141 genes, with two large clusters containing 44(C) and 57(E) genes, respectively. Cluster C was shared among a majority of the corn *Xvv* strains, and some genes are also present in *Xvh*. All genes from Cluster E are shared by *Xvh* and U.S. and Argentinian *Xvv* strains, while absent of most South African *Xvv* strains (Figure 3).

**Figure 4.**
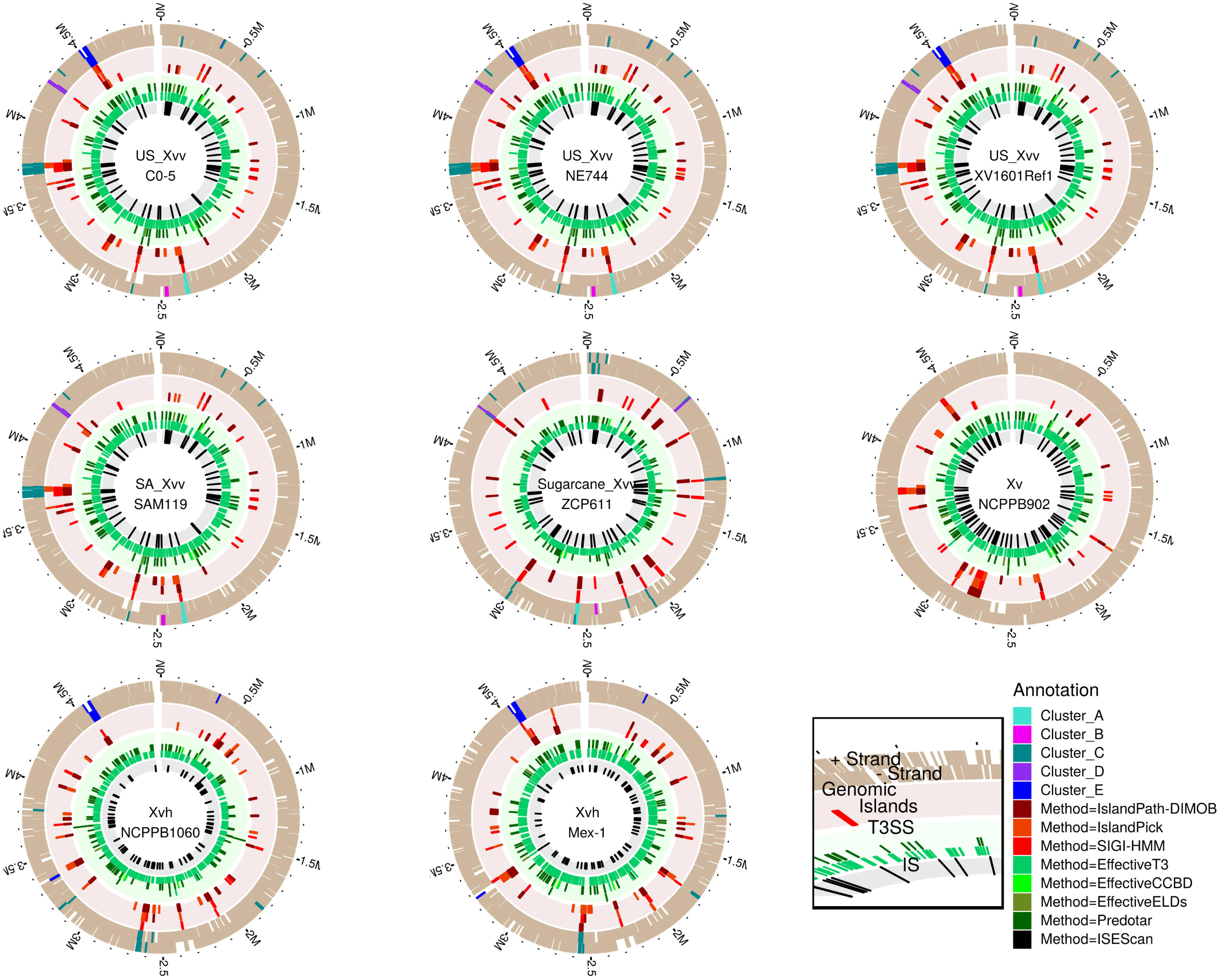
Genomic islands, predicted type 3 effectors and insertion sequences in *X. vasicola* genomes. Circular representation of eight complete *X. vasicola* genomes with annotated regions of interest. Legend in black square at the bottom right indicates what each circle represents. Genomic scale in million base pairs is shown. The outermost circles show presence of annotated genes in each genome. Clusters of genes identified as over-represented in US-*Xvv*, and their orthologues, are shown in colors. Predicted Genomic Islands using three methods integrated in Island Viewer (Bertelli et al. 2017) are shown in red colors, multiple predictions for a same region are shown stacked, SIGI-HMM is based on sequence composition, IslandPath-DIMOB is based on dinucleotide bias and presence of mobility genes, and IslandPick is based on phylogenetic comparisons. Predictions of type 3 secreted proteins are shown in green colors, four methods integrated in the EffectiveDB (Eichinger et al. 2016) suite are shown: EffectiveT3 predicts Type 3 secretion signals, Effective CCBD, conserved binding domains of Type 3 chaperones, EffectiveELD, eukaryotic-like domains, and Predotar predicts plant subcellular localization. Proteins having a significant score with at least EffectiveT3 are shown. Insertion sequence (IS) elements are shown in black as identified using ISEScan (Xie and Tang 2017).

The annotations and predicted functions of over-represented genes showed an enrichment in proteins with nucleic binding activity and involvement in DNA metabolism, recombination and transposition, suggesting mobile elements in the strains (Supplementary Figure 5). No particular enrichment was found in the group of underrepresented genes.

**Figure 5.**
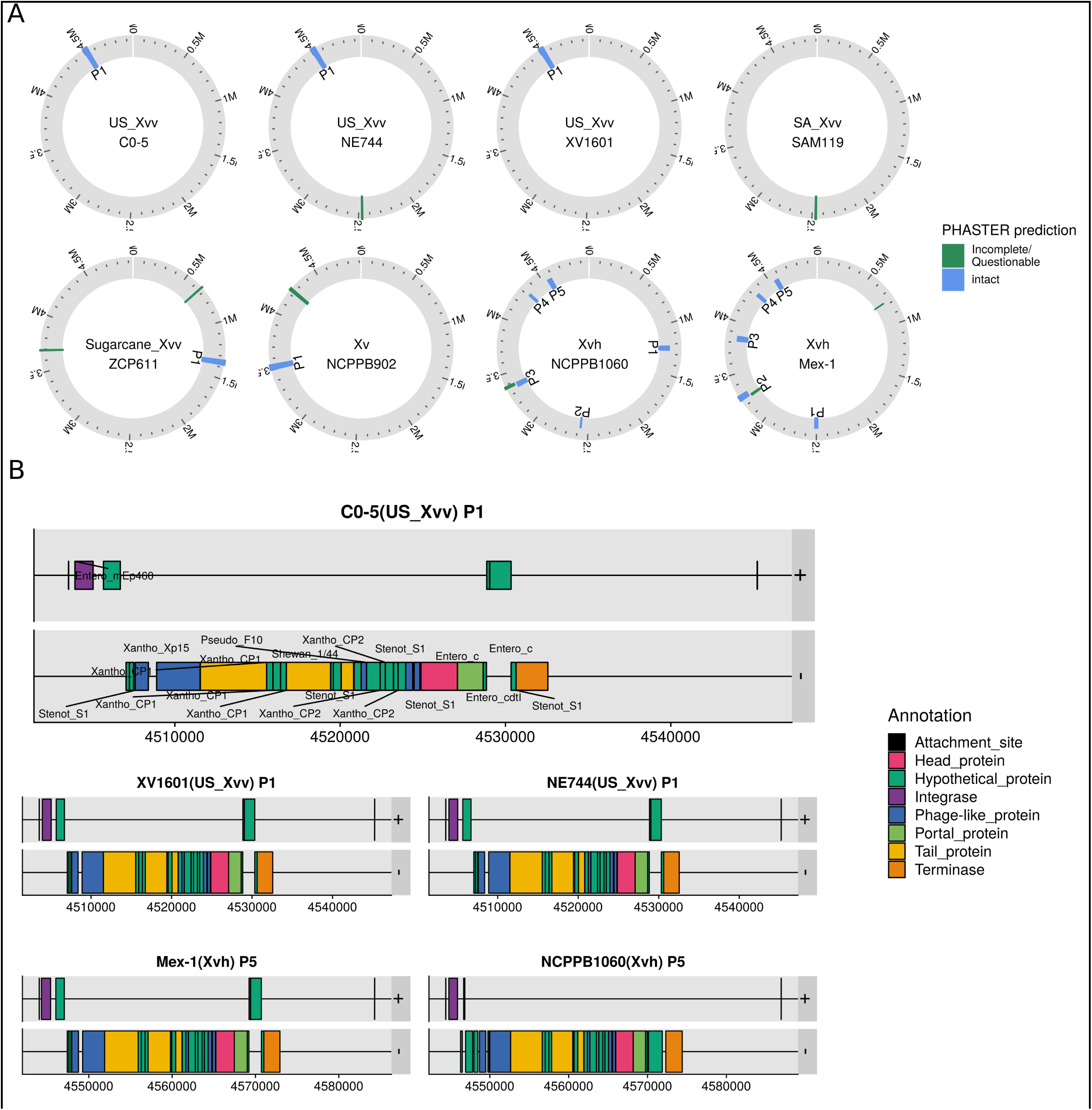
Prophages in *X. vasicola* genomes. **A).** Circular representation of identified prophages in eight complete *X. vasicola* genomes, genomic scale in million base pairs is shown. Intact prophage regions as identified by PHASTER (Arndt et al. 2016) are shown in blue and named P+number according to their position. Incomplete or questionable regions are shown in green (regions that are close together are shown stacked to improve readability). B) Diagram showing the genes in the predicted prophage corresponding to Cluster E in genomes of *Xvh* and *Xvv.* The genes are grouped according to their strand and colored according to their annotation. The diagram for *Xvv* strain CO-5 P1 shows for each gene the top hit against known genes in the Virus and Prophage database of PHASTER (Arndt et al. 2016), genes with no annotation had no significant hits.

### Clusters of genes in U.S. *Xvv* are genomic islands and contain putative effectors

We further analyzed these clusters in a set of eight fully sequenced genomes. Most of the identified clusters (except Cluster B) are predicted to be in genomic islands, using the IslandViewer4suite, which compiles parametric and phylogenetic methods for genomic island prediction (Bertelli et al. 2017) and implies that the clusters were acquired by horizontal transfer (Figure 4). Furthermore, Cluster C is particularly enriched in insertion sequences (IS) transposition-associated genes (Figure 4 and Supplementary Figure 5). IS were absent in Cluster E.

None of the over or underrepresented genes matched against known Type III effectors (T3E) by blast (Altschul et al. 1997). Furthermore, no specific association was found between effector presence/absence and U.S. *Xvv* or corn *Xvv*, in general, with the possible exception of XopG1, a M27 zinc protease (White et al. 2009), which is absent in most *Xvv* strains and present in other *X. vasicola* pathovars (Supplementary Figure 6).

**Figure 6.**
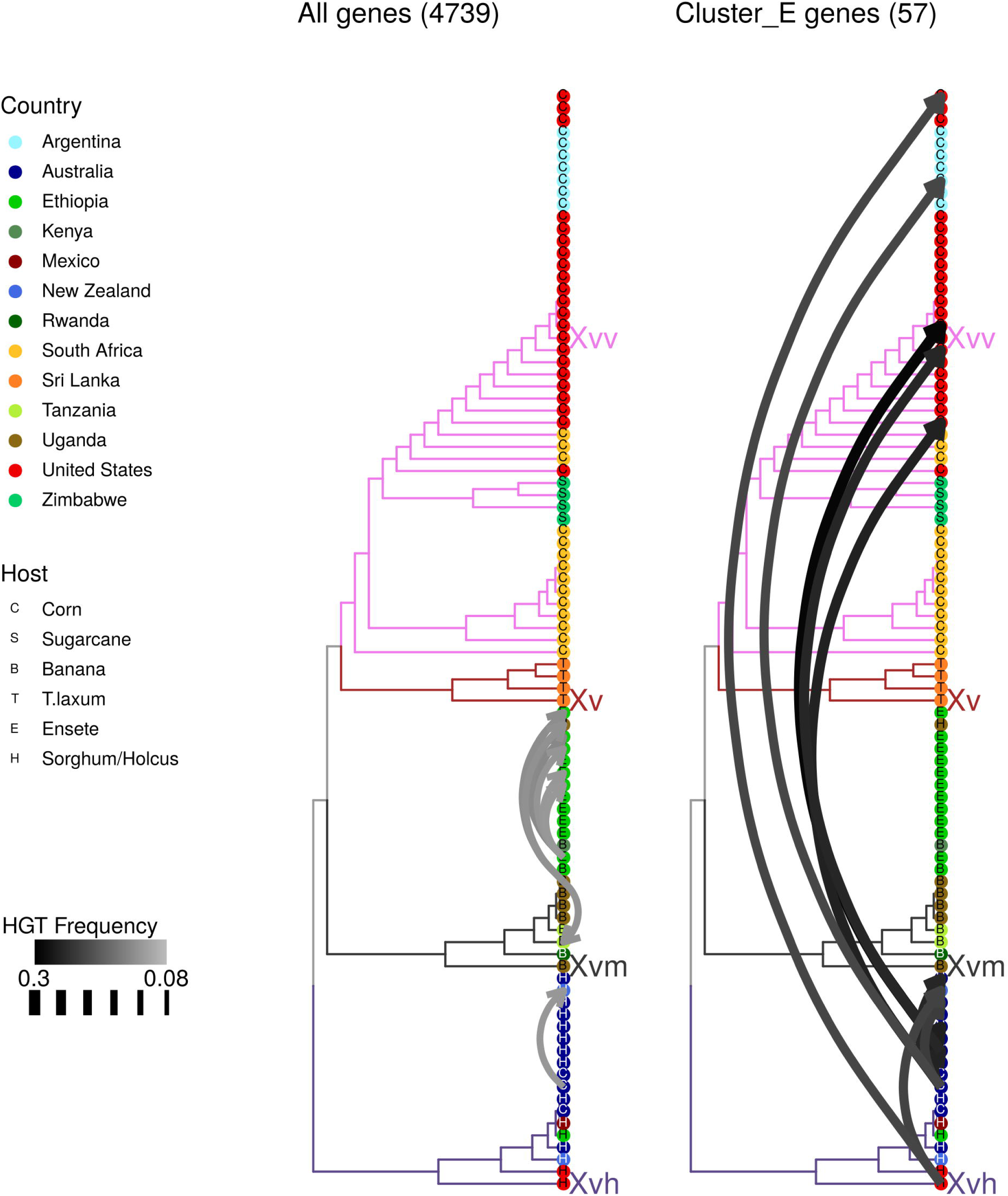
Horizontal Gene Transfer events predicted between *X. vasicola* genomes. A strain tree based on all core genome orthologs and ortholog gene trees obtained with orthofinder (Emms and Kelly 2015) were reconciled using Ranger-DTL (Bansal et al. 2018) to identify possible horizontal gene transfer events. Tips of the trees indicate country and host of isolation for each genome and branch colors indicate the four main *X. vasicola* pathovars. Tree to the left shows the results for all ortholog trees, and to the right for genes assigned to U.S. *Xvv* cluster E. Arrow thickness and color indicate predicted cumulative frequency of each event, a frequency of one would mean an event was identified in all 100 evaluated reconciled trees for all genes analyzed. Top 10 events with highest probability for each set are shown. Arrow heads indicate the direction of the predicted event.

Additionally, the suite EffectiveDB, which allows identification of putative T3Es based on prediction of secretion signals, T3 chaperone binding domains, eukaryotic-like domains and eukaryotic subcellular localization (Eichinger et al. 2016), predicted that ∼450 proteins were putative T3Es in each genome (having, at minimum, a predicted T3E secretion signal) (Figure 4). Some predicted T3Es proteins were found in the clusters, including five genes in Cluster C and nine in Cluster E, the latter included various hypothetical proteins, an HTH transcriptional regulator, and two methyltransferases (Supplementary Table 3).

### A gene cluster in U.S. *Xvv* is a prophage horizontally transferred from *Xvh*

Since these clusters are likely to have been horizontally transferred we attempted to find the taxonomic origin of the transfer by using Kaiju (Menzel, Ng, and Krogh 2016) to find the closest match for each gene from the CO-5 strains in the progenomes database (Mende et al. 2017), a database of representative microbial genomes that does not include *Xvv* or *Xvh*. As a whole, over 93% of the CO-5 genome was effectively assigned to *Xanthomonas* sp., with most genes assigned to *Xvm* (Supplementary Figure 7). In contrast, the gene clusters contained sequences from different taxonomic groups as well as several unassigned sequences. In Cluster A, eight genes (67%), were assigned to *Pantoea ananatis* (Supplementary Figure 7), which is frequently isolated along with *Xvv* (Lang et al. 2017; Coutinho et al. 2015). Within Cluster C, 40% of the genes were assigned to groups other than *Xanthomonas* including species of *Sphingobium* and *Pseudomonas* (Supplementary Figure 7).

Meanwhile in Cluster E, 57% of the genes were assigned to *Xanthomonas* sp., but curiously, 8% of the genes matched phages in the Caudovirales group (Supplementary Figure 7). This cluster was also enriched in GO terms associated to viral life cycles (Supplementary Figure 5), and unlike the other clusters, was not associated to insertion sequences (Figure 4), so its acquisition could have been mediated by phage transmission.

The finding of phage-related sequences prompted us to scan the genomes for additional phage sequences using PHASTER (Arndt et al. 2016) Indeed Cluster E corresponded to an intact prophage that included a region larger than 30 kb containing a high percentage of phage-related proteins. This was the only prophage identified in U.S. *Xvv* strains (Figure 5). Both *Xvh* strains examined contained the prophage in a similar genomic location, but also contained four other intact prophages. No prophage was identified in the South African *Xvv* strains, and strains from sugarcane and *T. laxum* both contained a prophage different from the one corresponding to Cluster E (Supplementary Figure 8). Many of the proteins in the Cluster E prophage have similarities to proteins from the *Xanthomonas*-infecting phages Cp1 and Cp2 (Figure 5).

The Cluster E/prophage genes are found in all U.S. and Argentinian corn *Xvv*, and some genes are found in one contemporary South African strain (Xvz45) (Figure 3). Older South African strains do not contain prophage genes, and neither do *Xv, Xvm* or sugarcane *Xvv.* On the other hand, all *Xvh* isolates contain these genes despite when isolated. A scenario that could explain this distribution is that this region was acquired at some point in *Xvh* and was recently horizontally transferred to an ancestor of the Argentinian and U.S. *Xvv* populations.

To explore this scenario we used Ranger-DTL (Bansal et al. 2018) to reconcile a whole genome phylogenetic tree with gene trees for each ortholog group. In each reconciliation, the more likely horizontal gene transfer (HGT) events and their direction were identified. When analyzing all suitable ortholog trees (4739 in total), various possible exchanges where identified, mostly within pathovars, and more abundantly within *Xvm* (Figure 6). As for genes in Cluster E, the results suggested that indeed, a transfer from *Xvh* to the U.S. and Argentina *Xvv* clade occurred. However, since many of the genes in this Cluster are identical across strains (Supplementary Figure 9), it was not possible to establish a clear origin or destination of the transfer. Similar results were obtained for Cluster E using ALE (amalgamated likelihood estimation), another reconciliation technique, albeit with lower probabilities (Supplementary Figure 10).

Overall, our results identified possible regions associated with the emerging *Xvv* population infecting corn in the U.S. and Argentina. These regions potentially contain virulence determinants or genes that conferred an advantage to this population for corn colonization in a way that explains its rapid proliferation.

## Discussion

In this work we show that the emerging populations of *Xvv* infecting corn in Argentina and the U.S. are genetically related and have acquired genomic regions, specifically a prophage (Cluster E) that may be associated with their spread. In phylogenetic analyses, the Argentinian *Xvv* strains are closer to South African strains than U.S. strains, suggesting a possible South American origin for the current epidemic. Accordingly, Argentinian *Xvv* strains also appear connected to *Xvh* strains indicating the horizontal gene transfer of the prophage from *Xvh* to *Xvv* could have occurred in South America (Figure 1). One U.S. strain (NE-7), grouped closer to Argentinian strains, which is consistent with at least two separate introductions to the U.S. Alternative scenarios are also possible since at least one contemporaneous South African *Xvv* strain (Xvz45) contained some of the genes in the cluster and another strain (XvzGP) grouped with U.S. strains.

The rapid spread of the disease in the U.S. and the possible ongoing genetic exchange between these distant populations may have been facilitated by human activity. Corn breeders in the U.S. accelerate product development by maintaining year-long operations and have increasingly adopted the practice of using winter nurseries for breeding and seed production (Butruille et al. 2015; Brewbaker 2003); many of these winter nurseries are located in the southern hemisphere, including South America (Zaworski 2016; Butruille et al. 2015) Additionally, corn production and export in South America has experienced considerable growth in the last decades (Meade et al. 2016). Although it is still unknown whether *Xvv* is transmitted by seeds, current practices likely allow enough exchange of contaminated material such that an adapted population can spread quickly between continents.

When inoculated on corn, Argentinian and U.S. *Xvv* strains caused more severe symptoms than South African strains, indicating that the emerging American populations are phenotypically different than the older (1988) South African population. We were unable to test contemporary South African strains because none are available in collections. Testing of newer populations will be needed to determine if the current disease spread is associated with an increase in virulence, since it is also possible that the tested South African strains have reduced virulence due to their extended time in storage. Testing different corn accessions will also be needed to confirm a gain of virulence since phenotypes seem to vary between varieties (Lang et al. 2017).

We identified five clusters of genes that were over-represented in the U.S. *Xvv* strains. These gene clusters overlapped with predicted genomic island regions, consistent with acquisition through horizontal gene transfer. Several clusters may represent important genomic acquisitions, if not for the current emerging population, for corn *Xvv* strains overall. Cluster C, for instance, is a group of ∼44 genes found in all contemporary corn *Xvv* strains (although not all strains contain all genes) and some *Xvh* strains. This cluster is enriched in mobility genes: transposases, insertion sequences, and various DNA binding genes, and it contains 5 predicted T3 secreted proteins. Taxonomic analyses revealed that a large percentage of genes in this cluster match other groups of bacteria including *Pseudomonas*, S*phingobium* and various *Burkholderiales.*

Cluster A contained eight genes found in *Pantoea ananatis*, including genes involved in replication (replication proteins A and C) and conjugation (P-type conjugative transfer proteins). *P. ananatis* is the causal agent of brown stalk rot of corn (Goszczynska et al. 2007) but it is also a versatile organism able to infect monocotyledonous and dicotyledonous hosts, and it is also a common epiphyte and endophyte (Coutinho and Venter 2009). *P. ananatis* was documented in association with *Xvv* on *Eucalyptus* in S. Africa (Coutinho et al. 2015) and with *Xvv* BLS symptomatic corn in the U.S. (Lang et al. 2017). However, the *Pantoea* strains alone were unable to cause BLS symptoms in corn (Lang et al. 2017), and brown stalk rot symptoms have not been reported on plants infected with BLS. The relationship between *Xvv* and *P. ananatis* is intriguing and it is possible *Xvv* may have acquired important virulence capacity from this association.

We focused on Cluster E because it is associated with the Argentina and U.S. *Xvv* populations, and thus may be related to their emergence. The prophage region in Cluster E is shared by *Xvh* and U.S. and Argentinian *Xvv*, and contained genes resembling elements of *Xanthomonas-*infecting bacteriophages CP1, CP2 and Xp10. This prophage is absent in other groups, including a recently reclassified available genome of a strain isolated from Areca nut (Wicker et al, this issue) (Bradbury 1986) that is closely related to *Xvv* (absence of the prophage was verified using PHASTER (Arndt et al. 2016)).

Prophages are temperate (non-infective or non-lytic) viruses that are integrated into bacterial genomes by recombination (Varani et al. 2013). They are important vehicles for horizontal gene transfer, they can promote recombination and rearrangements in the bacterial genome, and they often carry additional non-essential cargo genes (morons) that may confer new phenotypic properties to the bacteria (Varani et al. 2013; Brüssow, Canchaya, and Hardt 2004). Prophages have been known to carry virulence factors or factors that enhance bacterial fitness (Brüssow, Canchaya, and Hardt 2004; Figueroa-Bossi et al. 2001), although reduction of virulence has also been reported (Ahmad et al. 2014). Prophages harboring elements conferring virulence activity have been found in different plant pathogenic bacteria including *Xylella* sp. *(Varani et al. 2008)* and *Candidatus* Liberibacter asiaticus (Jain, Fleites, and Gabriel 2015). And in *Xanthomonas arboricla*, strains pathogenic on walnut carry a higher number (and also a different repertoire) of prophages than non-pathogenic strains (Cesbron et al. 2015).

We hypothesize that the Cluster E prophage region contains genes that play a role in virulence in *Xvh*, and when horizontally transferred to *Xvv*, it enhanced virulence or fitness to the emerging *Xvv* populations. In *Xvh*, Cluster E was found in all examined strains, and was one of only two shared pro-phages between two geographically and temporarily distant *Xvh* strains analyzed (1961 Zimbabwe vs 2016 U.S.) (Figure 5, Supplementary Figure 8). Furthermore most of the genes in this cluster were identical across all compared *Xvv* and *Xvh* (Supplementary Figure 9), suggesting they are not subject to prophage decay (Brüssow, Canchaya, and Hardt 2004) and may indeed play a beneficial role for the bacteria. This cluster contains genes predicted to be T3-secreted, peptidases and transcription factors that could have virulence activity, and various non-phage related hypothetical proteins with unknown function. Further characterization of these genes, as well as of other over-represented *Xvv* genes that were not assigned to clusters, is needed to establish a possible role in virulence.

Here we have used comparative genomics to address questions about the origin of an epidemic and the genetic determinants associated with pathogen population spread. Based on our findings, we postulate different exciting hypotheses that will be the subject of future work to understand the lifestyle and evolution of *Xvv* and related bacteria.

## Materials and Methods

### Strain collection and molecular detection

Isolation of *Xvv* from corn leaves was performed as in Lang et al. (2017) with minor modifications. Instead of placing the tissue in distilled water, fresh tissues were dissolved in 1 mL of 10 mM MgCl_2_, macerated with sterile pellet pestle and incubated for at least 1.5 hours at room temperature. For bacterial isolation, one loop-fill (10 µL) of solution was spread onto nutrient agar (NA). Plates were incubated at 28°C for two days. Single characteristic bright yellow colonies were selected, and re-streaked for further isolation until pure colonies were obtained. Samples from the United States were collected across several fields in Colorado, Iowa, Kansas and Nebraska. Samples from Argentina were collected from fields located in San Luis, Córdoba, and Santa Fe states. (Supplementary Table 1).

South African *Xvv* strains where obtained from the L. E. Claflin collection (Qhobela, Claflin, and Nowell 1990). Australian *Xvh* strains were obtained from the NSW Department of Primary Industries Plant Pathology and Mycology Herbarium Culture Collection (https://www.dpi.nsw.gov.au/about-us/services/collections/herbarium).

Molecular detection of *Xvv* was performed following one of these procedures: For the cases using colony PCR, one single colony was suspended in 10 µL of sterile water and boiled at 95°C for 5 min. First, colony PCR of suspected *Xvv* samples were performed using Xvv3 or Xvv5 primers as described previously (Lang et al. 2017). To further confirm isolates, a second method using 16S rRNA gene and a housekeeping gene, atpD (ATP synthase β chain), was used to identify bacteria to species level. PCR reactions for 16S rRNA (50 µL) contained 2 µL of boiled DNA template, 0.2 µM of each primer (Supplementary Table 4), 1X GoTaq reaction buffer, 2 mM MgCl2, 0.2 mM dNTP, and 0.25 unit/µL GoTaq DNA polymerase enzyme (Promega, Madison, WI). The cycling conditions were as follows: initial denaturation at 94°C for 3 min, following 35 cycles of 94°C for 45 s, 50°C for 1 min and 72°C for 1:30 min, and the final extension period at 72°C for 10 min. PCR fragments were separated in a 1.5% agarose gel for 45 min at 90 V, and fragments were extracted and purified using the DNA clean & concentrator kit (ZYMO Research). Sequencing was performed with 5 ng/µL of each PCR product at Quintara Biosciences (Fort Collins, CO) and analyzed using Geneious software (version 10.0.7). Sequence identities to the genus level were determined using Blastn from the NCBI database.

### Disease phenotyping on corn

Corn (hybrid P1151) was grown in a 1:1 mix of Promix-BX Biofungicide + Mycorrhizae (Quakertown, PA) under greenhouse conditions (30 ± 1 °C, 16 hour day length, and 80% relative humidity). Three weeks after planting, plants were inoculated with 24 *Xvv* isolates and one *Xvh* isolate (Supplementary Table 1). Each bacterial strain was cultured in peptone sucrose agar (PSA) for 24 hours at 28°C and then suspended to 10^8^ CFU mL^−1^ in sterile, distilled water. Bacterial suspensions were infiltrated as described by Lang et al. 2017. Two leaves were inoculated on at least seven individual plants. Infiltration experiments were repeated three times and data was combined to perform statistical analysis. Sterile, distilled water was used as a negative control in all inoculations. Quantification of the lesion length was done by measuring the expansion distance beyond the infiltration site at seven days post inoculation (dpi).

For statistical analysis a one-way ANOVA using lesion length ∼ Isolate was made using the aov function in R (R Core Team 2013), square root transformation of lesion length data was done to satisfy ANOVA requirements. Treatment groups were obtained using a Tukey’s HSD (honestly significant difference) test, with the HSD.test in the agricolae package (de Mendiburu and de Mendiburu 2019).

### Genome Sequencing, assembly and data collection

Genomic DNA for the *Xanthomonas* positive samples was extracted using Easy-DNA kit (Invitrogen) and PCR amplification of the *atpD* gene was carried out for further confirmation to the species level. PCR reactions for *atpD* gene (40 µL) contained 25 ng/µL of DNA template, 0.4 µM of each primer (Supplementary Table 4), 1X GoTaq reaction buffer, 1.5 mM MgCl2, 0.2 mM dNTPs, and 0.1 unit/µL GoTaq DNA polymerase enzyme (Promega, Madison, WI). Cycling conditions were performed as described by (Fargier, Saux, and Manceau 2011).

For 24 *Xvv* strains and one *Xvh* strain from the U.S., Illumina sequencing was performed by BGI (www.bgi.com) using HiSeq 4000 with paired-end 100 bp reads. All Illumina reads were first trimmed with Trimmomatic (PE ILLUMINACLIP:CO-2.adapters.fa LEADING:2 TRAILING:2 SLIDINGWINDOW:4:2 MINLEN:30) (Bolger, Lohse, and Usadel 2014) and then assembled into scaffolds using SPAdes (Bankevich et al. 2012) with default settings. For 11 Australian *Xvh* strains sequencing was performed using Illumina Miseq and assembled using the A5 pipeline (Coil, Jospin, and Darling 2015).

Five strains (CO-5, XV1601, NE744, Mex-1 and ZCP611) were sequenced using long read, single molecule real time sequencing (SMRT Sequel, PacBio, Menlo Park, CA). SMRT read sequences were assembled using HGAP v4 (Chin et al. 2013). Genomes were circularized using circlator (Hunt et al. 2015). For XV1601, Illumina reads were also available and were used to polish the PacBio assembly using Canu v1.3-r7616 (Koren et al. 2017). No major differences were found between with the polished assembly nor with the other SMRT-generated genomes as examined with multiple alignments with Mauve (Darling et al. 2004). All generated genomes have been deposited to the NCBI (Supplementary Table 1)

We obtained all available assemblies of *X. vasicola* (NCBI:txid56459) and *X. campestris* pv. *musacearum* (NCBI:txid454958, here referred to as *X. vasicola* pv. *musacearum*) as of November 2018 (Supplementary Table 1). Assemblies with an N50 of minimum 10 kbp were kept (thus excluding *Xvv* strains NCPPB895 and NCPPB890). For 10 recently published *Xvv* strains form South Africa (named *Xanthomonas vasicola* pv. *zeae)* (Sanko et al. 2018) the available assemblies ranged in size from 3.8 to 4.5 Mbp, significantly less than the average size for other *X. vasicola* genomes (∼4.9Mbp), and alignments to reference genomes revealed large fragments missing from the assemblies (Supplementary Figure 11). These genomes were thus reassembled from available Illumina raw reads (Biosample accessions SAMN10286417-26) using Unicycler v0.4.8-beta (--mode bold), which functions as a SPAdes-optimiser (Wick et al. 2017). New assemblies had expected sizes and were not missing large regions and were thus kept for analysis in this work (Supplementary Figure 11).

### Genome annotation and ortholog identification

All assemblies were automatically annotated using prokka v1.14-dev (--rfam) (Seemann 2014). Ortholog groups from prokka-annotated proteins were identified using Orthofinder v. 2.2.6 (default parameters) (Emms and Kelly 2015). Additionally, orthologs were also identified, and core genome size was estimated, using Pan-x (Ding, Baumdicker, and Neher 2018) Similar ortholog groups with similar distribution were found with both strategies (Pan-X = 6155 groups and 2163 unassigned genes, Orthofinder = 6084 groups and 1896 unassigned genes), since Orthofinder grouped more genes together, these results were kept for further analyses.

### Phylogenetic analyses

Phylogenetic trees were used were obtained with various methods using whole genome data. KSNP3 (Gardner, Slezak, and Hall 2015) was used to obtain parsimony and maximum-likelihood trees based on pan-genome SNPs from identified k-mers (K-mer size = 21), the parsimony tree had overall higher branch support and was kept for analysis. CSI Phylogeny 1.4 (Kaas et al. 2014) was used to obtain trees based on core genome SNPs, with *Xoo* PXO99A used as reference for SNP calling. Trees based on whole genome protein alignments obtained using the STAG method implemented in orthofinder (Emms and Kelly 2015) and RAxML+FasTree in Pan-X (Ding, Baumdicker, and Neher 2018; Price, Dehal, and Arkin 2010; Stamatakis 2014) were also analyzed.

MLSA neighbor-joining trees were obtained by identifying 31 housekeeping genes using AMPHORA v2 (Kerepesi, Bánky, and Grolmusz 2014), creating multiple alignments form their concatenated sequences using MUSCLE v3.8.31 (Edgar 2004), and generating the trees using functions of the R package phangorn (pml, optim.pml (model = “Blosum62”) and bootstrap.pml(bs=100)) (Schliep 2011). Average nucleotide identity values were obtained using the ANI-matrix script from the enveomics collection (v1.3) (Rodriguez-R and Konstantinidis 2016).

Minimum spanning trees were generated with MSTGold v2.5 (Salipante and Hall 2011) using a multiple alignment of core genome SNPs identified with CSI Phylogeny (Kaas et al. 2014), a consensus tree of eight estimated different MSTs out of maximum 3000 tested was kept and bootstraped 500 times (parameters = -n 3000 -m 43200 -b 10 -t 50 -s 500).

RangerDTL was used to explore reconciliations between species trees and gene trees for all identified orthologs using the dated method (Bansal et al. 2018). Gene and species trees used were generated by orthofinder (Emms and Kelly 2015); the species tree was made ultrametric for this analysis using the chronos function of the ape package ((Paradis, Claude, and Strimmer 2004). For each gene, 100 trees (using variable –seed from 1 to 100) were reconciled with the species tree and possible horizontal gene transfer (HGT) were identified with a probability corresponding to the number of trees were the event was identified. To analyze multiple genes simultaneously (as for Cluster E), the probabilities for each event in each tree were averaged. Amalgamated likelihood estimation, ALE v0.5 (Szöllősi Gergely J. et al. 2015) was also used to perform the same analysis with the same trees.

### Identification of over-represented regions

A hypergeometric test was designed and applied to each ortholog group identified with orthofinder to look for over or under-represented genes in corn U.S. *Xvv* strain. The test was applied using the function phyper(q, m, n, k, lower.tail = TRUE, log.p = FALSE) in R, where for each ortholog: q = strains in the U.S. *Xvv* group that contain the gene, m=total number of strains in the U.S. *Xvv* group, n=number of strains in the comparison group, k=total strains that contain the gene in both groups.

The test was applied for each gene in both directions, for over-representation (q – 1) and under-representation, and the lowest *p* value was chosen (if the lowest *p* value was for the under-representation test, it was multiplied by −1 to differentiate them). The test was applied 100 times for each gene, each time changing the comparison group by randomly selecting a group of non-U.S. *Xvv* strains of a random size between 10 and 69 (total of non-U.S. *Xvv* strains) (Supplementary Figure 4). The average *p* value of the 100 tests was taken, and a correction for multiple testing (p.adjust function, method BH (Benjamini and Hochberg 1995) in R) was applied to the *p* values obtained for all genes.

Genes with an absolute adjusted *p* value < 0.05 were considered as over or under-represented in the U.S. *Xvv* group. The position of the selected genes in the genome of the strain CO-5 was then used to establish clusters. Groups of more than 10 genes over-represented genes found less than 5 kb from each other were considered a Cluster and assigned a letter (A-E) according to their distance to the replication origin.

### Annotation of genomic regions

For a more thorough annotation of genes in each clusters and to assess enrichment in functional categories, protein sequences of the CO-5 strain were further annotated using Blast2GO v5.2.5 (Conesa et al. 2005) by combining hits against the ncbi nr-protein database (blast-p fast, e-value 0.01, number of hits 10), InterPro, Gene Onthology terms (GOs), and KEGG enzyme codes (default parameters). Enrichment of GO terms was assessed for the different groups using a hyper-geometric test as implemented in the GoFuncR package (Grote 2018).

Genomic Islands were predicted using the IslandViewer 4 suite (Bertelli et al. 2017) Insertion Sequences (IS) were identified using ISEScan (v1.6) (default parameters) (Xie and Tang 2017). And possible prophage were identified using PHASTER (Arndt et al. 2016). All three analyses were made using prokka-annotated files for strains with complete genomes.

Known Type III (T3) effectors were identified by blastp (v. 2.6.0+, results were filtered keeping hits with -evalue < 0.0001, >30% identity in >40% the query length) (Altschul et al. 1997) of consensus effectors sequences obtained from http://xanthomonas.org/ against the protein sequences obtained using Prokka. Novel T3 effectors were predicted using effective DB (default parameters + plant model for Predotar) (Eichinger et al. 2016), results were filtered to keep proteins with an EffectiveT3 (signal peptide) of minimum 0.9999, plus any additional predictions with other methods.

The web version of Kaiju (Menzel, Ng, and Krogh 2016) was used to annotate possible taxonomic origin of cluster genes against the progenomes database (default parameters) (Mende et al. 2017).

### Visualization and other analyses

Most figures were generated using R (R Core Team 2013). Phylogenetic trees were generated using the ggtree package (Yu et al. 2017). Circular genome and genomic region visualizations were generated using ggbio (Yin, Cook, and Lawrence 2012). Heatmaps were generated using pheatmap (Kolde and Kolde 2015). Upset plot was generated using UpsetR (Conway, Lex, and Gehlenborg 2017). Comparisons of genomic regions were made using GenomicRanges (Lawrence et al. 2013). Genomic alignments were visualized using Mauve v Jan-19-2018 (Darling et al. 2004).

## Acknowledgments

We would like to thank collaborators Tamra Jackson-Ziems, Terra Hartman, Silvina Areas and Garry Munkvold for providing isolates of Xvv from Nebraska and Iowa. This work was funded by grants from the Colorado Corn Administrative Committee, APHIS (project # 6.0533.01) and the Foundation for Food and Agriculture Research (project # 544722)

## Supplementary Materials Legends

**Supplementary Figure 1. Phylogeny of *X. vasicola* strains obtained using different methods.** Trees are shown using different methods: Orthofinder and Pan-X build trees based on protein sequences of core genome trees using the STAG method and RaxML+FasTree respectively. CSI phylogeny builds trees based on core genome SNPs, and the MLSA tree was generated based on concatenated sequences of housekeeping genes identified with AMPHORA, aligned with Muscle. The tree was built by using the R package phangorn. When present, the trees are rooted using *X. oryzae* pv. *oryzae* (*Xoo*) PXO99A as an outgroup, otherwise the tree was rooted using *Xvh* NCPPB 1060. *Xoo* PXO99A was excluded from pan-X analyses. Bars to the left indicate branch length as generated by each program.

**Supplementary Figure 2. Average Nucleotide Identity (ANI) between pairs of *X. vasicola* genomes.** The heatmap shows pairwise ANI values between *X. vasicola* genomes, with *Xoo* PXO99A included for comparison. Dendrograms to the top and left show hierarchical clustering of genomes based on ANI. Bars to the left indicate Host, Pathovar and Country of isolation.

**Supplementary Figure 3. Shared orthologs between different *X. vasicola* groups. A)** UpSet visualization of intersections between orthologs present in each relevant *X. vasicola* group. Orthologs were identified using orthofinder in each genome, an ortholog group was said to be present in a group if it was present in at least 30% of the strains evaluated. Vertical bars show the intersection between groups with bold circles below. First bar corresponds to the intersection of all groups (core genome). Horizontal bars indicate the number of genes found in each group. Highlighted in blue is the intersection between corn *Xvv* from the U.S. and Argentina with *Xvh*, and highlighted in purple are genes exclusive to corn *Xvv.* B) Core genome statistics obtained from Pan-X. Percentage of core and accessory genes is shown (left), then number of strains containing groups of orthologs (middle) and the distribution of gene length in all genomes (right).

**Supplementary Figure 4. Identification of over-represented genes in U.S. Xvv.** A) The frequency of each gene (each point) in the U.S. *Xv* population (x axis) is compared to their frequency in sets of randomly chosen *X. vasicola* genomes (y axis). The average frequency of each gene in 100 groups is shown and error bars indicate standard deviation. Dot colors indicate whether a given gene was identified as over or under-represented. B) Density plot showing uniform size distribution for random sets of genomes (100 per gene) chosen as comparison groups for hypergeometric tests to determine over or under representation when compared to U.S. *Xvv* genomes C) Density plot shows the pathovar composition of the random sets.

**Supplementary Figure 5. Gene Ontology (GO) term enrichment in over-represented U.S. *Xvv* genes.** Go terms identified as statistically enriched in the group of over-represented genes and their genomic clusters are shown. No terms were found enriched for Clusters A, B or D. Dot color and size indicate *p* value of enrichment as determined using GoFuncR. GO annotations were obtained using Blast2GO.

**Supplementary Figure 6. Known Type 3 effectors in *X. vasicola* genomes.** Heatmap shows copy number of known T3 effectors as determined by blast of each genome against consensus *Xanthomonas* T3 effector sequences. Copy number is shown to a maximum of 3. The only effector with a higher copy number is AvrBs3 (TAL effectors) in *Xoo* PXO99A. Dendrogram at the top corresponds to the KSNP3 tree in Figure 1. Dendrogram to the left shows hierarchical clustering of effectors according to their presence/absence pattern in the genomes. Color bars at the top indicate Pathovar, Host and Country of isolation for each genome.

**Supplementary Figure 7. Taxonomic distribution of over-represented genes in possible horizontally transferred clusters in U.S. *Xvv.*** Krona plots obtained using Kaiju showing the taxonomic assignation of genes in each over-represented U.S. *Xvv* cluster in the strain CO-5 as well as in the whole genome. Each gene was matched to its closest sequence in the progenomes database and assigned a taxonomic group accordingly. Colors in each plot are ordered according to percentages and do not correspond to the same taxa across clusters.

**Supplementary Figure 8. Other prophages identified in *X. vasicola* genomes.** Diagrams show genes found in the predicted prophages in *X. vasicola* genomes different from the prophage corresponding to Cluster E. Gare grouped according to their strand and colored according to their annotation in phaster. For each gene the top hit against known genes in the Virus and Prophage database of PHASTER is shown; genes with no annotation had no significant hits.

**Supplementary Figure 9. Genetic distances of genes in ortholog groups assigned to over-represented clusters in U.S. *Xvv.*** Phylogenetic trees obtained with orthofinder were analyzed to find the distances between all tips in the tree (strains containing each gene) using the cophenetic function from the ape package. Boxplots show the distribution of distances for each tree, boxplots showing means around zero indicate that all the tips were found at the same distance, meaning the gene sequence was identical across strains.

**Supplementary Figure 10. Horizontal Gene Transfer events predicted between *X. vasicola* genomes using ALE.** A strain tree based on all core genome orthologs and ortholog gene trees obtained with orthofinder were reconciled using ALE to identify possible horizontal gene transfer events. Tips of the trees indicate country and host of isolation for each genome and branch colors indicate the four main *X. vasicola* pathovars. Tree to the left shows the results for all ortholog trees, and to the right for genes assigned to U.S. *Xvv* cluster E. Arrow thickness and color indicate predicted cumulative frequency of each event, a frequency of one would mean an event was identified in all 100 evaluated reconciled trees for all genes analyzed. Arrow head indicate the direction of the predicted event.

**Supplementary Figure 11. Reassembly of Xvv strains from South Africa.** Mauve multiple alignment shows the south African *Xvv* strain SAM119 (top), the publicly available assembly of strain Xvz45 (GCF_003111905.1) (middle), and a reassembly of strain Xvz45 used in this paper using the corresponding raw reads (SAMN10286417) (bottom). Long vertical red lines indicate contig limits. Similar results were obtained with other genomes form this set, indicated with accession numbers SAMN-in Supplementary Table 1.

**Supplementary Table 1. Inventory of genomic sequences used in this work.**

**Supplementary Table 2. Ortholog groups determined as over or under-represented in U.S. *Xvv* strains**

**Supplementary Table 3. Annotation of genes assigned to over-represented clusters in *Xvv* strain CO-5.**

**Supplementary Table 4. Primers used for molecular detection of *Xvv.***

## References

Ahmad, A. A., Askora, A., Kawasaki, T., Fujie, M., and Yamada, T. 2014. The filamentous phage XacF1 causes loss of virulence in Xanthomonas axonopodis pv. citri, the causative agent of citrus canker disease. Front Microbiol. 5:321.

Altschul, S. F., Madden, T. L., Schäffer, A. A., Zhang, J., Zhang, Z., Miller, W., et al. 1997. Gapped BLAST and PSI-BLAST: a new generation of protein database search programs. Nucleic Acids Res. 25:3389–3402.

Arndt, D., Grant, J. R., Marcu, A., Sajed, T., Pon, A., Liang, Y., et al. 2016. PHASTER: a better, faster version of the PHAST phage search tool. Nucleic Acids Res. 44:W16–W21.

Bankevich, A., Nurk, S., Antipov, D., Gurevich, A. A., Dvorkin, M., Kulikov, A. S., et al. 2012. SPAdes: A New Genome Assembly Algorithm and Its Applications to Single-Cell Sequencing. Journal of Computational Biology. 19:455–477.

Bansal, M. S., Kellis, M., Kordi, M., and Kundu, S. 2018. RANGER-DTL 2.0: rigorous reconstruction of gene-family evolution by duplication, transfer and loss. Bioinformatics. 34:3214–3216.

Benjamini, Y., and Hochberg, Y. 1995. Controlling the false discovery rate: a practical and powerful approach to multiple testing. Journal of the Royal statistical society: series B (Methodological). 57:289–300.

Bertelli, C., Laird, M. R., Williams, K. P., Lau, B. Y., Hoad, G., Winsor, G. L., et al. 2017. IslandViewer 4: expanded prediction of genomic islands for larger-scale datasets. Nucleic Acids Res. 45:W30–W35.

Bolger, A. M., Lohse, M., and Usadel, B. 2014. Trimmomatic: a flexible trimmer for Illumina sequence data. Bioinformatics. 30:2114–2120.

Bradbury, J. F. 1986. Guide to plant pathogenic bacteria. CAB international.

Brewbaker, J. L. 2003. Corn production in the tropics: The Hawaii experience. University of Hawaii.

Broders, K. 2017. Status of bacterial leaf streak of corn in the United States. In Proceedings of the Integrated Crop Management Conference, Iowa State University, Digital Press. Available at: https://lib.dr.iastate.edu/icm/2017/proceedings/18/ [Accessed February 25, 2019].

Brüssow, H., Canchaya, C., and Hardt, W.-D. 2004. Phages and the Evolution of Bacterial Pathogens: from Genomic Rearrangements to Lysogenic Conversion. Microbiol Mol Biol Rev. 68:560–602.

Butruille, D. V., Birru, F. H., Boerboom, M. L., Cargill, E. J., Davis, D. A., Dhungana, P., et al. 2015. Maize Breeding in the United States: Views from Within Monsanto. In Plant Breeding Reviews: Volume 39, John Wiley & Sons, Ltd, p. 199–282.

Cesbron, S., Briand, M., Essakhi, S., Gironde, S., Boureau, T., Manceau, C., et al. 2015. Comparative Genomics of Pathogenic and Nonpathogenic Strains of Xanthomonas arboricola Unveil Molecular and Evolutionary Events Linked to Pathoadaptation. Front Plant Sci. 6:1126.

Chin, C.-S., Alexander, D. H., Marks, P., Klammer, A. A., Drake, J., Heiner, C., et al. 2013. Nonhybrid, finished microbial genome assemblies from long-read SMRT sequencing data. Nature Methods. 10:563–569.

Coil, D., Jospin, G., and Darling, A. E. 2015. A5-miseq: an updated pipeline to assemble microbial genomes from Illumina MiSeq data. Bioinformatics. 31:587–589.

Conesa, A., Götz, S., García-Gómez, J. M., Terol, J., Talón, M., and Robles, M. 2005. Blast2GO: a universal tool for annotation, visualization and analysis in functional genomics research. Bioinformatics. 21:3674–3676.

Conway, J. R., Lex, A., and Gehlenborg, N. 2017. UpSetR: an R package for the visualization of intersecting sets and their properties. Bioinformatics. 33:2938–2940.

Coutinho, T. A., and Wallis, F. M. 1991. Bacterial Streak Disease of Maize (Zea mays L.) in South Africa. Journal of Phytopathology. 133:112–112.

Coutinho, T. A., and Venter, S. N. 2009. Pantoea ananatis: an unconventional plant pathogen. Molecular Plant Pathology. 10:325–335.

Coutinho, T. A., Westhuizen, L. van der Roux, J., McFarlane, S. A., and Venter, S. N. 2015. Significant host jump of Xanthomonas vasicola from sugarcane to a Eucalyptus grandis clone in South Africa. Plant Pathology. 64:576–581.

Darling, A. C. E., Mau, B., Blattner, F. R., and Perna, N. T. 2004. Mauve: Multiple Alignment of Conserved Genomic Sequence With Rearrangements. Genome Res. 14:1394–1403.

Ding, W., Baumdicker, F., and Neher, R. A. 2018. panX: pan-genome analysis and exploration. Nucleic Acids Res. 46:e5–e5.

Dyer, R. A. 1949. Botanical surveys and control of plant diseases. Farming in South Africa. 24:119–121.

Edgar, R. C. 2004. MUSCLE: multiple sequence alignment with high accuracy and high throughput. Nucleic Acids Res. 32:1792–1797.

Eichinger, V., Nussbaumer, T., Platzer, A., Jehl, M.-A., Arnold, R., and Rattei, T. 2016. EffectiveDB—updates and novel features for a better annotation of bacterial secreted proteins and Type III, IV, VI secretion systems. Nucleic Acids Res. 44:D669–D674.

Emms, D. M., and Kelly, S. 2015. OrthoFinder: solving fundamental biases in whole genome comparisons dramatically improves orthogroup inference accuracy. Genome Biology. 16:157.

Fargier, E., Saux, M. F.-L., and Manceau, C. 2011. A multilocus sequence analysis of Xanthomonas campestris reveals a complex structure within crucifer-attacking pathovars of this species. Systematic and Applied Microbiology. 34:156–165.

Figueroa-Bossi, N., Uzzau, S., Maloriol, D., and Bossi, L. 2001. Variable assortment of prophages provides a transferable repertoire of pathogenic determinants in Salmonella. Molecular Microbiology. 39:260–272.

Gardner, S. N., Slezak, T., and Hall, B. G. 2015. kSNP3.0: SNP detection and phylogenetic analysis of genomes without genome alignment or reference genome. Bioinformatics. 31:2877–2878.

Goszczynska, T., Botha, W. J., Venter, S. N., and Coutinho, T. A. 2007. Isolation and Identification of the Causal Agent of Brown Stalk Rot, A New Disease of Maize in South Africa. Plant Disease. 91:711–718.

Grote, S. 2018. GOfuncR: gene ontology enrichment using FUNC. R package version. 1.

Hunt, M., Silva, N. D., Otto, T. D., Parkhill, J., Keane, J. A., and Harris, S. R. 2015. Circlator: automated circularization of genome assemblies using long sequencing reads. Genome Biology. 16:294.

Jain, M., Fleites, L. A., and Gabriel, D. W. 2015. Prophage-Encoded Peroxidase in ‘Candidatus Liberibacter asiaticus’ Is a Secreted Effector That Suppresses Plant Defenses. MPMI. 28:1330–1337.

Kaas, R. S., Leekitcharoenphon, P., Aarestrup, F. M., and Lund, O. 2014. Solving the Problem of Comparing Whole Bacterial Genomes across Different Sequencing Platforms. PLOS ONE. 9:e104984.

Kerepesi, C., Bánky, D., and Grolmusz, V. 2014. AmphoraNet: the webserver implementation of the AMPHORA2 metagenomic workflow suite. Gene. 533:538–540.

Kolde, R., and Kolde, M. R. 2015. Package ‘pheatmap.’ R Package. 1.

Koren, S., Walenz, B. P., Berlin, K., Miller, J. R., Bergman, N. H., and Phillippy, A. M. 2017. Canu: scalable and accurate long-read assembly via adaptive k-mer weighting and repeat separation. Genome Res. 27:722–736.

Korus, K., Lang, J. M., Adesemoye, A. O., Block, C. C., Pal, N., Leach, J. E., et al. 2017. First Report of Xanthomonas vasicola Causing Bacterial Leaf Streak on Corn in the United States. Plant Disease. 101:1030.

Lang, J. M., DuCharme, E., Ibarra Caballero, J., Luna, E., Hartman, T., Ortiz-Castro, M., et al. 2017. Detection and Characterization of Xanthomonas vasicola pv. vasculorum (Cobb 1894) comb. nov. Causing Bacterial Leaf Streak of Corn in the United States. Phytopathology. 107:1312–1321.

Lawrence, M., Huber, W., Pages, H., Aboyoun, P., Carlson, M., Gentleman, R., et al. 2013. Software for computing and annotating genomic ranges. PLoS computational biology. 9:e1003118.

Leite, R. P., Custódio, A. a. P., Madalosso, T., Robaina, R. R., Duin, I. M., and Sugahara, V. H. 2019. First Report of the Occurrence of Bacterial Leaf Streak of Corn Caused by Xanthomonas vasicola pv. vasculorum in Brazil. Plant Disease. 103:145–145.

Meade, B., Puricelli, E., McBride, W. D., Valdes, C., Hoffman, L., Foreman, L., et al. 2016. Corn and Soybean Production Costs and Export Competitiveness in Argentina, Brazil, and the United States. United States Department of Agriculture, Economic Research Service. Available at: https://ideas.repec.org/p/ags/uersib/262143.html.

Mende, D. R., Letunic, I., Huerta-Cepas, J., Li, S. S., Forslund, K., Sunagawa, S., et al. 2017. proGenomes: a resource for consistent functional and taxonomic annotations of prokaryotic genomes. Nucleic Acids Res. 45:D529–D534.

de Mendiburu, F., and de Mendiburu, M. F. 2019. Package ‘agricolae.’ R Package, Version.: 1.2-1.

Menzel, P., Ng, K. L., and Krogh, A. 2016. Fast and sensitive taxonomic classification for metagenomics with Kaiju. Nature Communications. 7:11257.

Moffett, M. L. 1983. Bacterial plant pathogens recorded in Australia. In Plant Bacterial Diseases: A Diagnostic Guide., Academic Press, Sydney., p. 317–336.

Paradis, E., Claude, J., and Strimmer, K. 2004. APE: Analyses of Phylogenetics and Evolution in R language. Bioinformatics. 20:289–290.

Péros, J. P., Girard, J. C., Lombard, H., Janse, J. D., and Berthier, Y. 1994. Variability of Xanthomonas Campestris pv. vasculorum From Sugarcane and Other Gramineae in Reunion Island. Characterization of a Different Xanthomonad. Journal of Phytopathology. 142:177– 188.

Plazas, M. C., De Rossi, R. L., Brücher, E., Guerra, F. A., Vilaró, M., Guerra, G. D., et al. 2017. First Report of Xanthomonas vasicola pv. vasculorum Causing Bacteria Leaf Streak of Maize (Zea mays) in Argentina. Plant Disease. 102:1026–1026.

Price, M. N., Dehal, P. S., and Arkin, A. P. 2010. FastTree 2 – Approximately Maximum-Likelihood Trees for Large Alignments. PLOS ONE. 5:e9490.

Qhobela, M., Claflin, L. E., and Nowell, D. C. 1990. Evidence that Xanthomonas campestris pv. zeae can be distinguished from other pathovars capable of infecting maize by restriction fragment length polymorphism of genomic DNA. Canadian Journal of Plant Pathology. 12:183–186.

R Core Team. 2013. R: A language and environment for statistical computing.

Rodriguez-R, L. M., and Konstantinidis, K. T. 2016. The enveomics collection: a toolbox for specialized analyses of microbial genomes and metagenomes. PeerJ Preprints. Available at: https://peerj.com/preprints/1900/ [Accessed February 7, 2017].

Salipante, S. J., and Hall, B. G. 2011. Inadequacies of Minimum Spanning Trees in Molecular Epidemiology. Journal of Clinical Microbiology. 49:3568–3575.

Sanko, T. J., Kraemer, A. S., Niemann, N., Gupta, A. K., Flett, B. C., Mienie, C., et al. 2018. Draft Genome Assemblages of 10 Xanthomonas vasicola pv. zeae Strains, Pathogens Causing Leaf Streak Disease of Maize in South Africa. Genome Announc. 6:e00532–18.

Schliep, K. P. 2011. phangorn: phylogenetic analysis in R. Bioinformatics. 27:592–593.

Seemann, T. 2014. Prokka: rapid prokaryotic genome annotation. Bioinformatics. 30:2068– 2069.

Stamatakis, A. 2014. RAxML version 8: a tool for phylogenetic analysis and post-analysis of large phylogenies. Bioinformatics. 30:1312–1313.

Szöllősi Gergely J., Davín Adrián Arellano, Tannier Eric, Daubin Vincent, and Boussau Bastien. 2015. Genome-scale phylogenetic analysis finds extensive gene transfer among fungi. Philosophical Transactions of the Royal Society B: Biological Sciences. 370:20140335.

Tushemereirwe, W., Kangire, A., Ssekiwoko, F., Offord, L. C., Crozier, J., Boa, E., et al. 2004. First report of Xanthomonas campestris pv. musacearum on banana in Uganda. Plant Pathology. 53:802–802.

USDA-NASS. 2017. Crop production Summary 2016, United States Department of Agriculture, National Agricultural Statistics Service. Washington, D.C.□: United States Department of Agriculture, Statistical Reporting Service, Crop Reporting Board□: [Supt. of Docs., U.S. G.P.O., distributor]. Available at: http://purl.access.gpo.gov/GPO/LPS1137.

Varani, A. de M., Souza, R. C., Nakaya, H. I., Lima, W. C. de, Almeida, L. G. P. de, Kitajima, E. W., et al. 2008. Origins of the Xylella fastidiosa Prophage-Like Regions and Their Impact in Genome Differentiation. PLOS ONE. 3:e4059.

Varani, A. M., Monteiro-Vitorello, C. B., Nakaya, H. I., and Van Sluys, M.-A. 2013. The Role of Prophage in Plant-Pathogenic Bacteria. Annual Review of Phytopathology. 51:429–451.

White, F. F., Potnis, N., Jones, J. B., and Koebnik, R. 2009. The type III effectors of Xanthomonas. Molecular Plant Pathology. 10:749–766.

Wick, R. R., Judd, L. M., Gorrie, C. L., and Holt, K. E. 2017. Unicycler: Resolving bacterial genome assemblies from short and long sequencing reads. PLOS Computational Biology. 13:e1005595.

Xie, Z., and Tang, H. 2017. ISEScan: automated identification of insertion sequence elements in prokaryotic genomes. Bioinformatics. 33:3340–3347.

Yin, T., Cook, D., and Lawrence, M. 2012. ggbio: an R package for extending the grammar of graphics for genomic data. Genome biology. 13:R77.

Young, J. M., Bradbury, J. F., Davis, R. E., Dickey, R. S., Ercolani, G. L., Hayward, A. C., et al. 1991. Nomenclatural revisions of plant pathogenic bacteria and list of names 1980-1988. Review of Plant Pathology. 70:211–221.

Yu, G., Smith, D. K., Zhu, H., Guan, Y., and Lam, T. T.-Y. 2017. ggtree: an R package for visualization and annotation of phylogenetic trees with their covariates and other associated data. Methods in Ecology and Evolution. 8:28–36.

Zaworski, F. 2016. Winter Breeding Programs in South America. SeedWorld. Available at: https://seedworld.com/winter-breeding-programs-south-america/ [Accessed February 21, 2019].

