## Supplementary figures and images for "Genomic acquisitions in emerging populations of *Xanthomonas vasicola* pv. *vasculorum* infecting corn in the U.S. and Argentina"

### Supplementary Figure 1

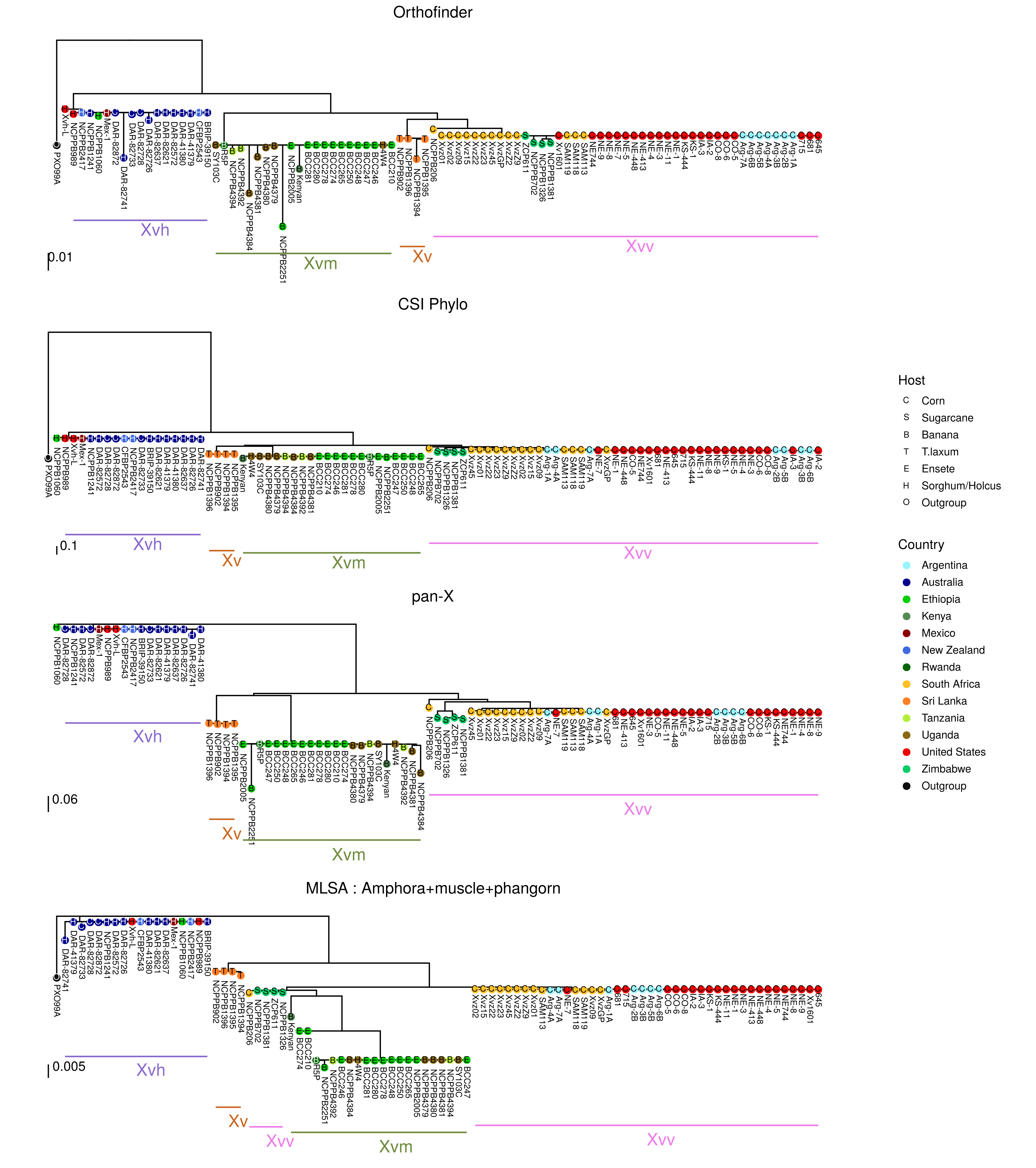

### Supplementary Figure 2

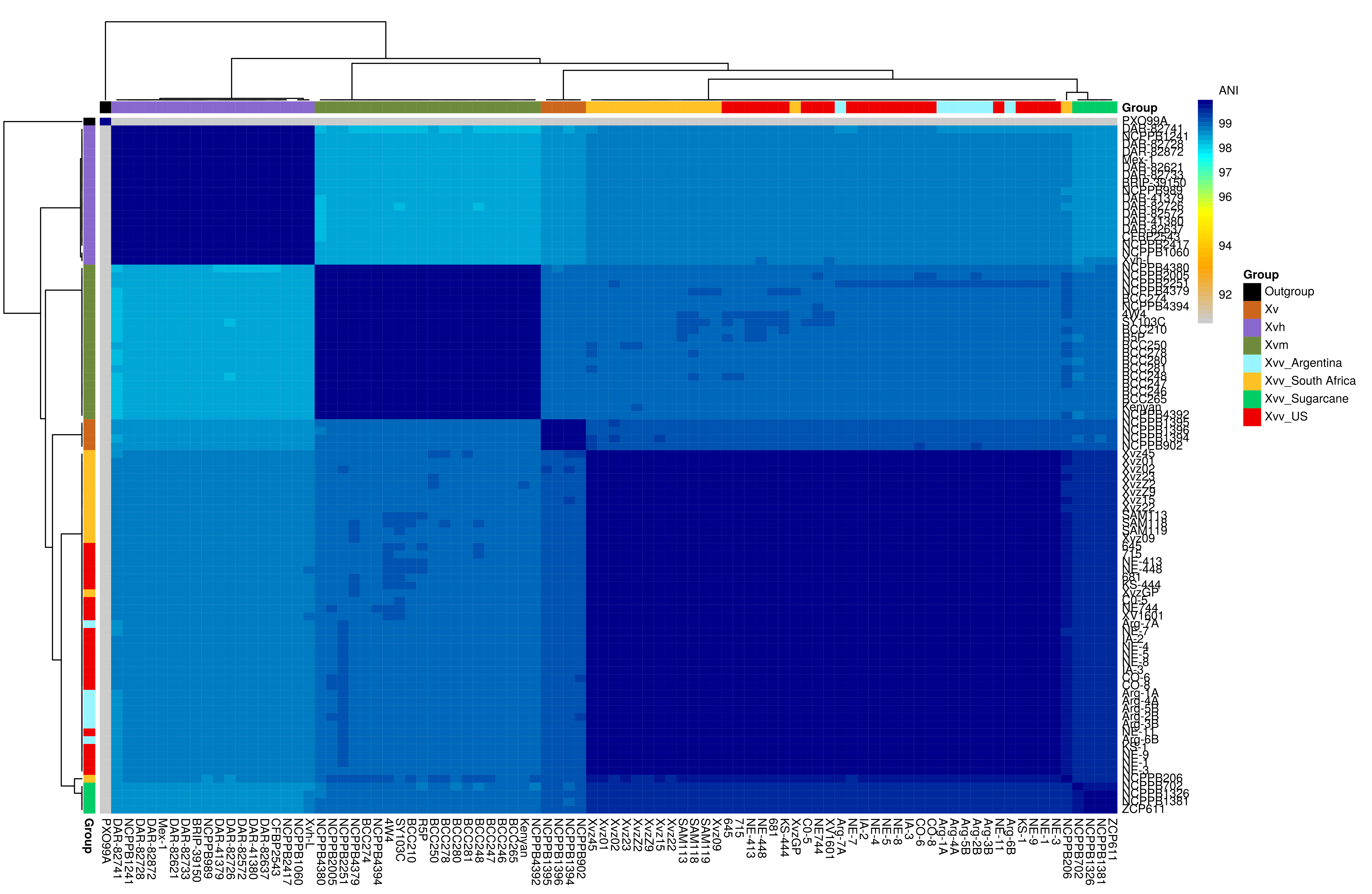

### Supplementary Figure 3

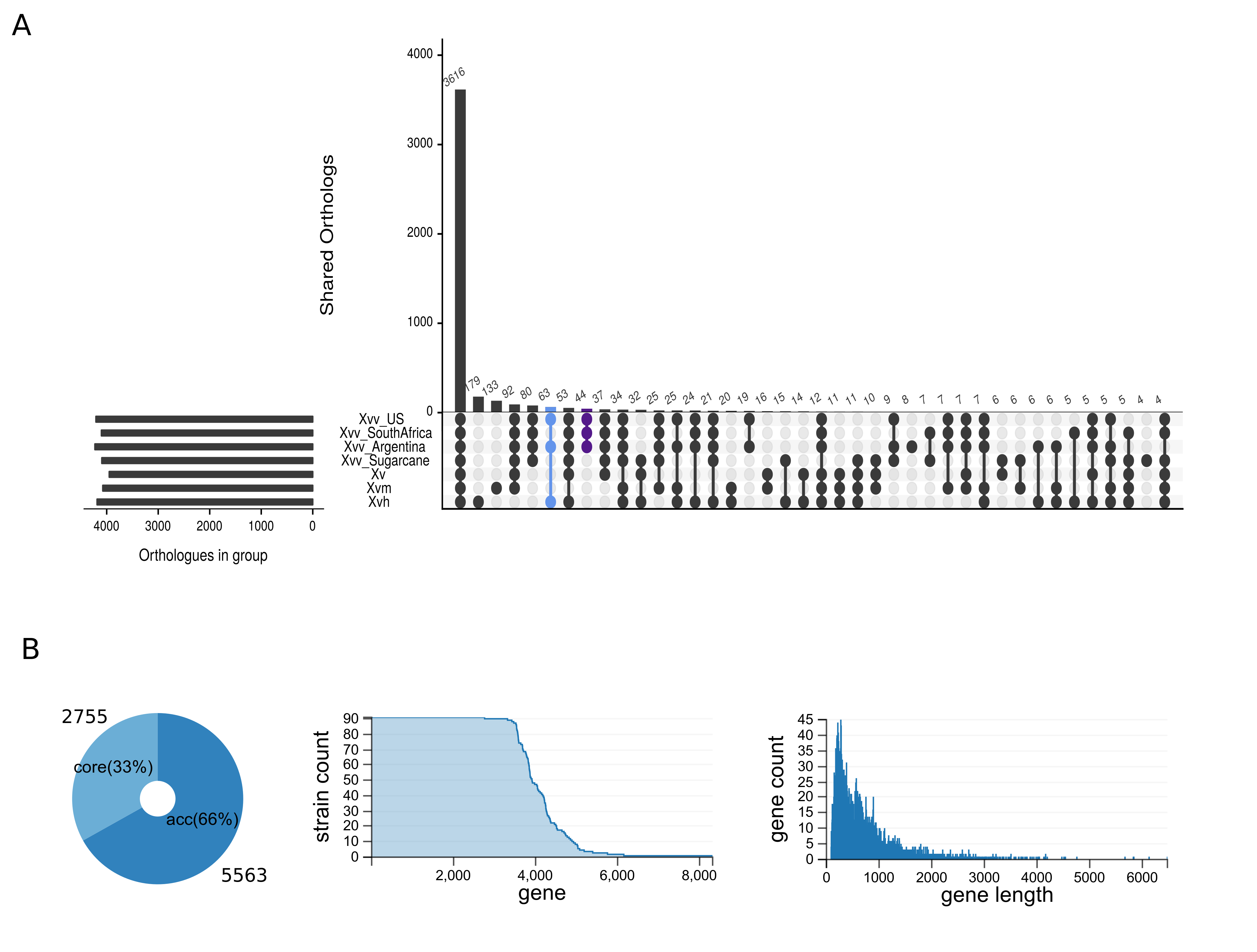

### Supplementary Figure 4

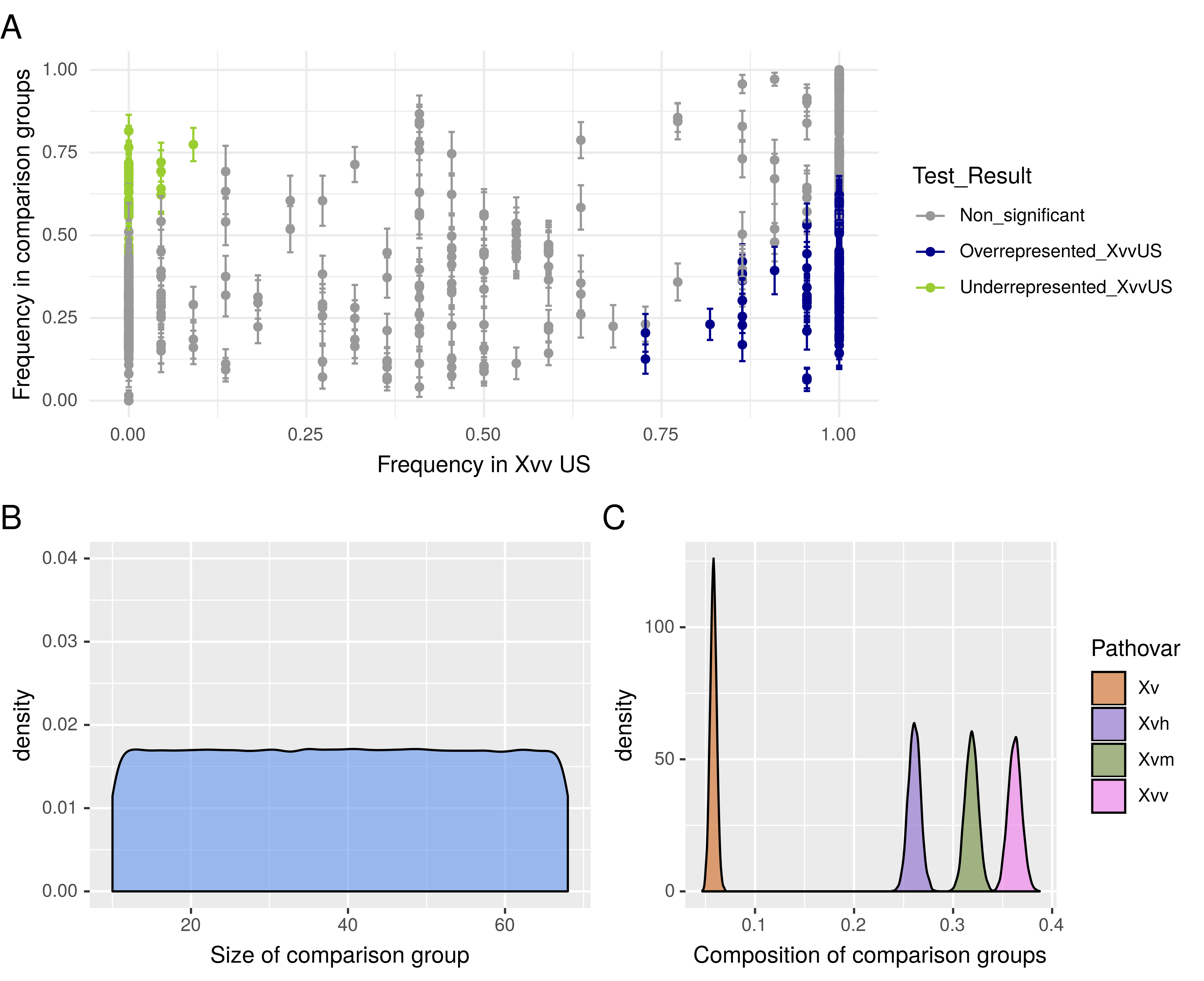

### Supplementary Figure 5

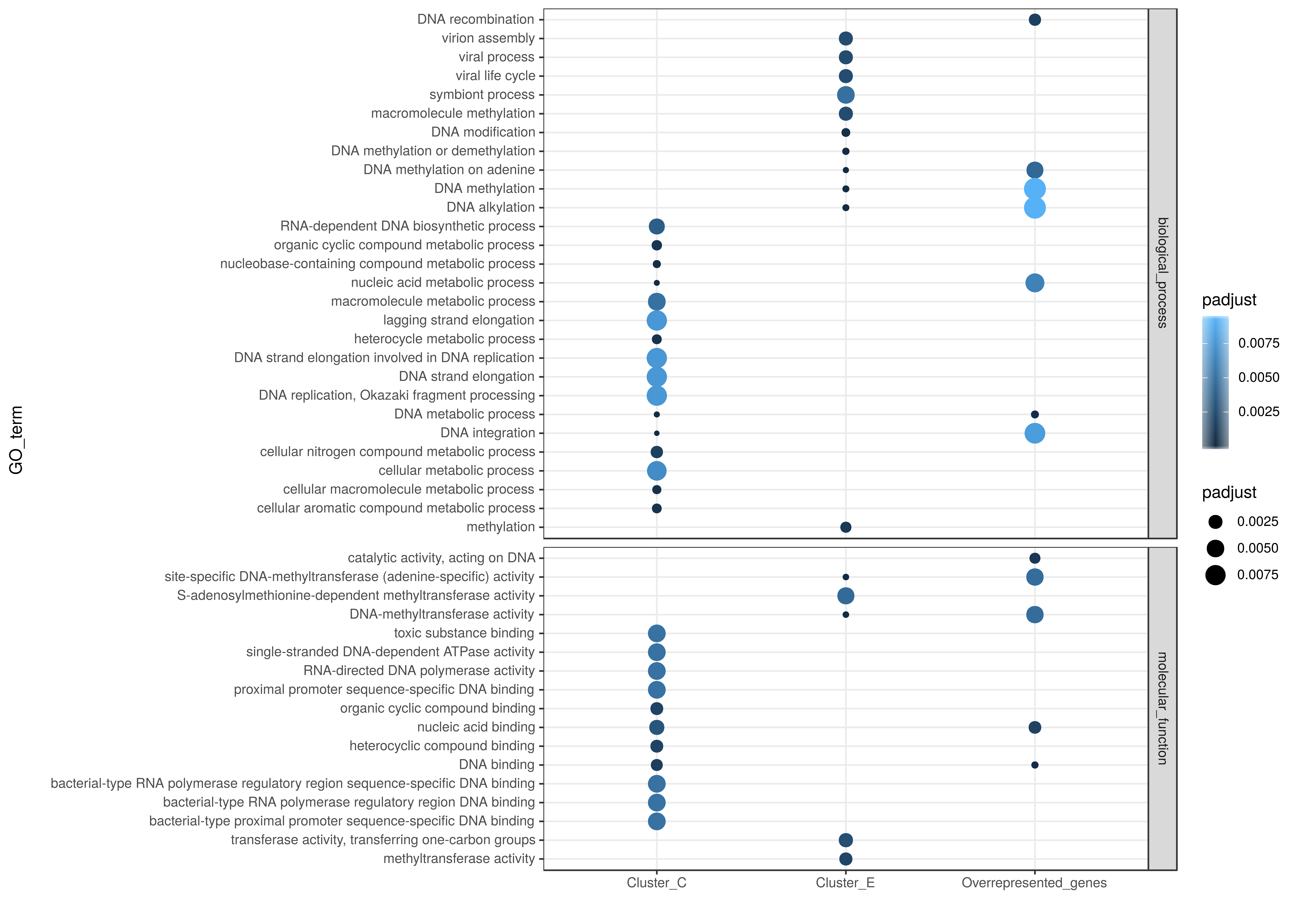

### Supplementary Figure 6

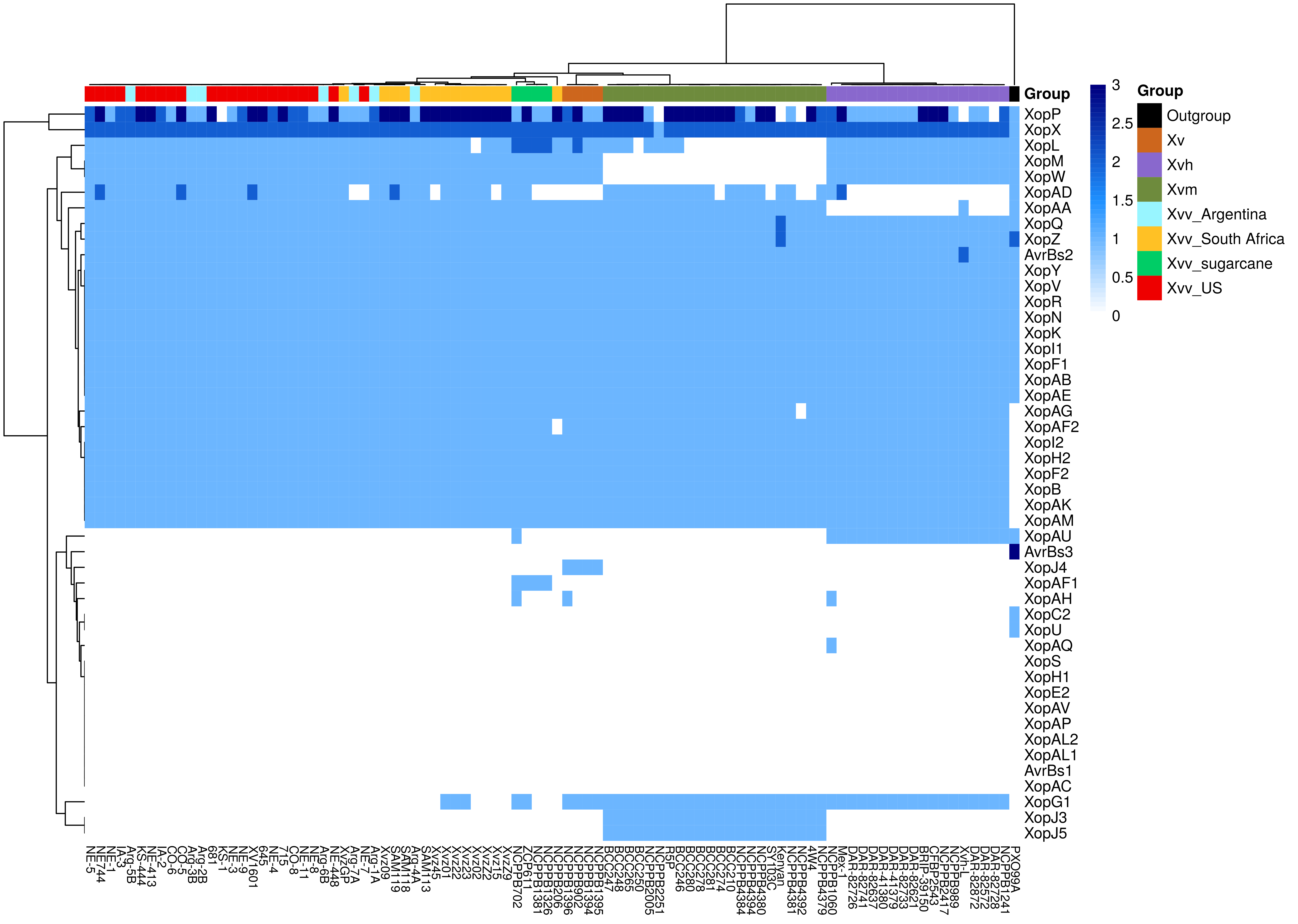

### Supplementary Figure 7

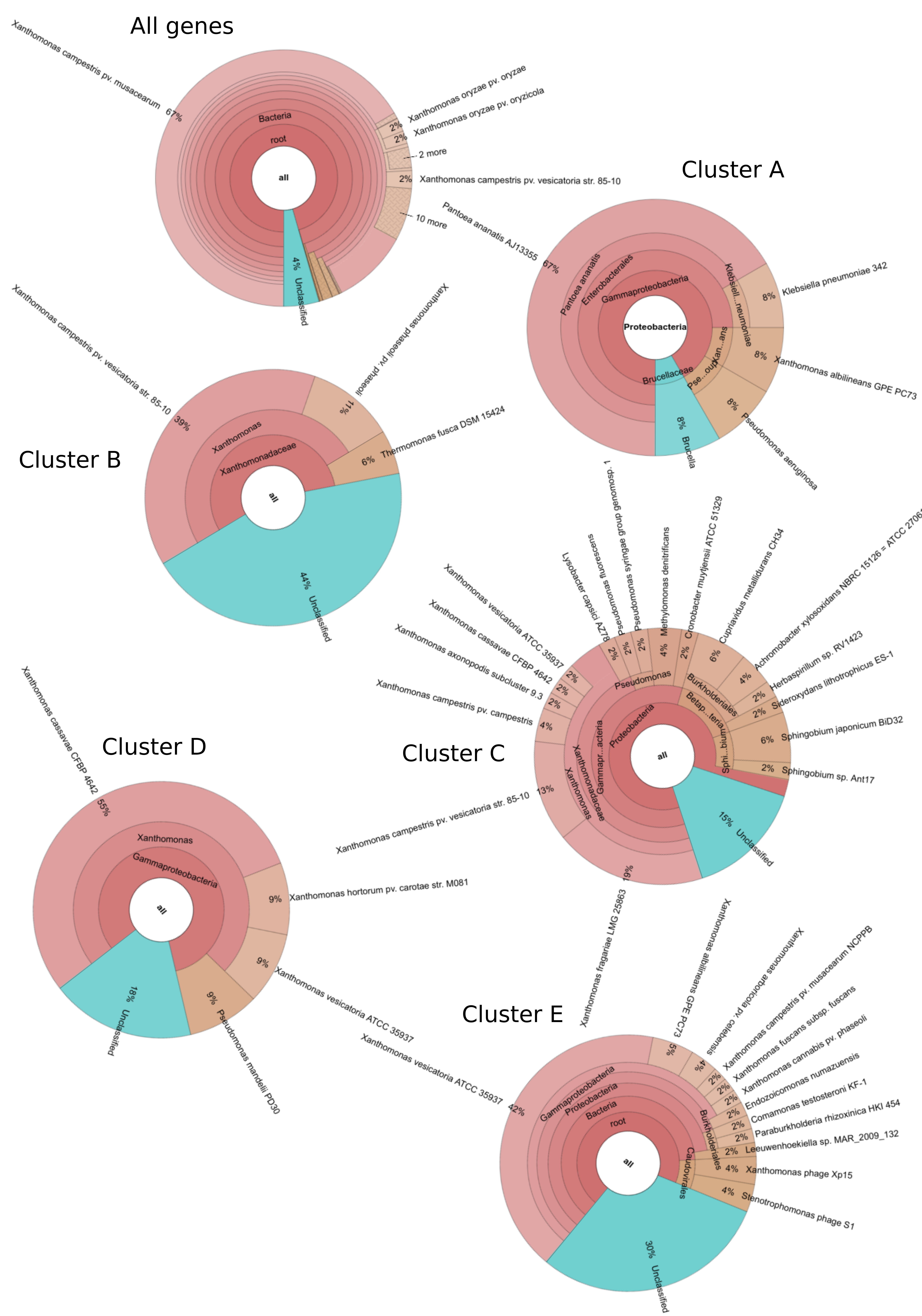

### Supplementary Figure 8

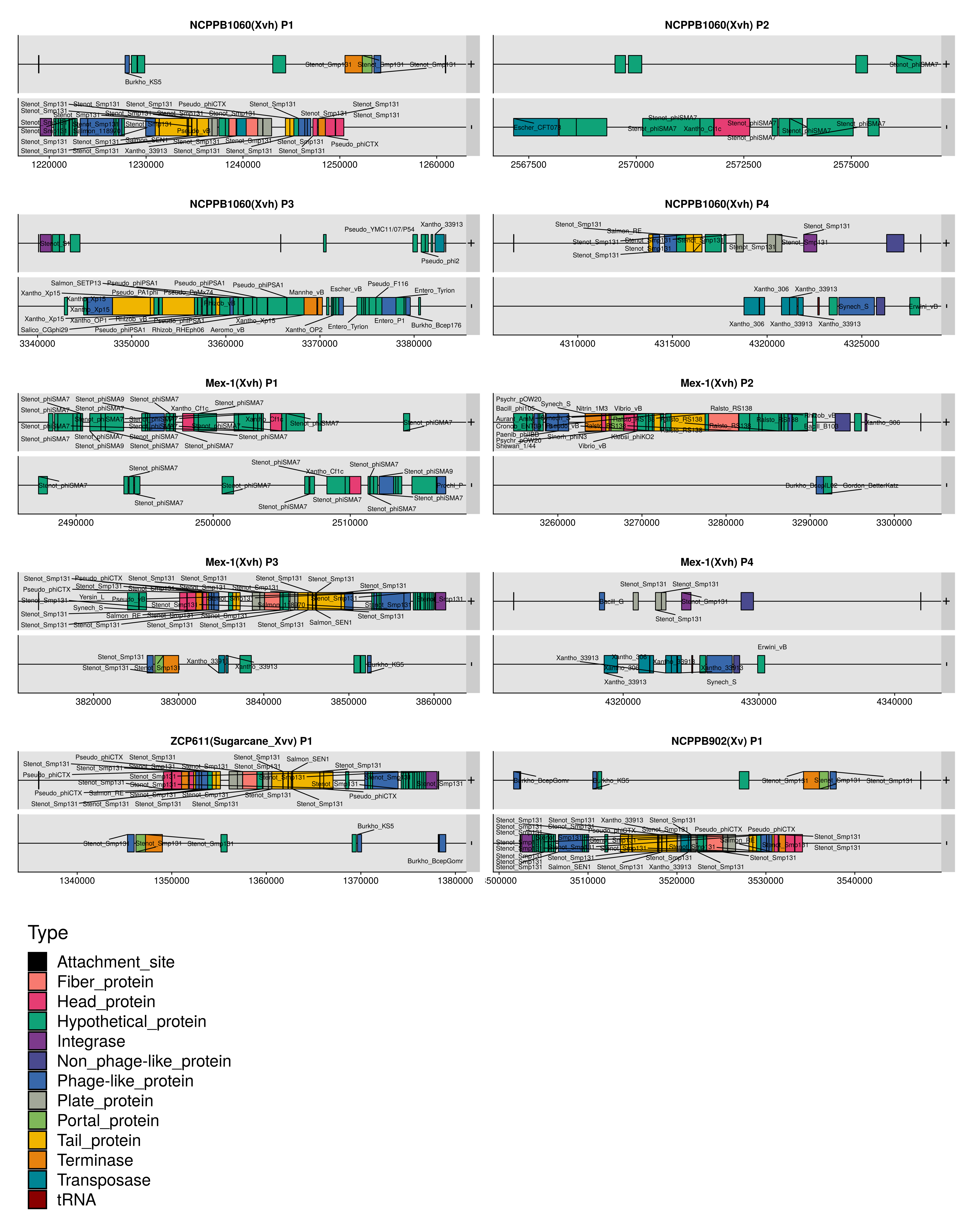

### Supplementary Figure 9

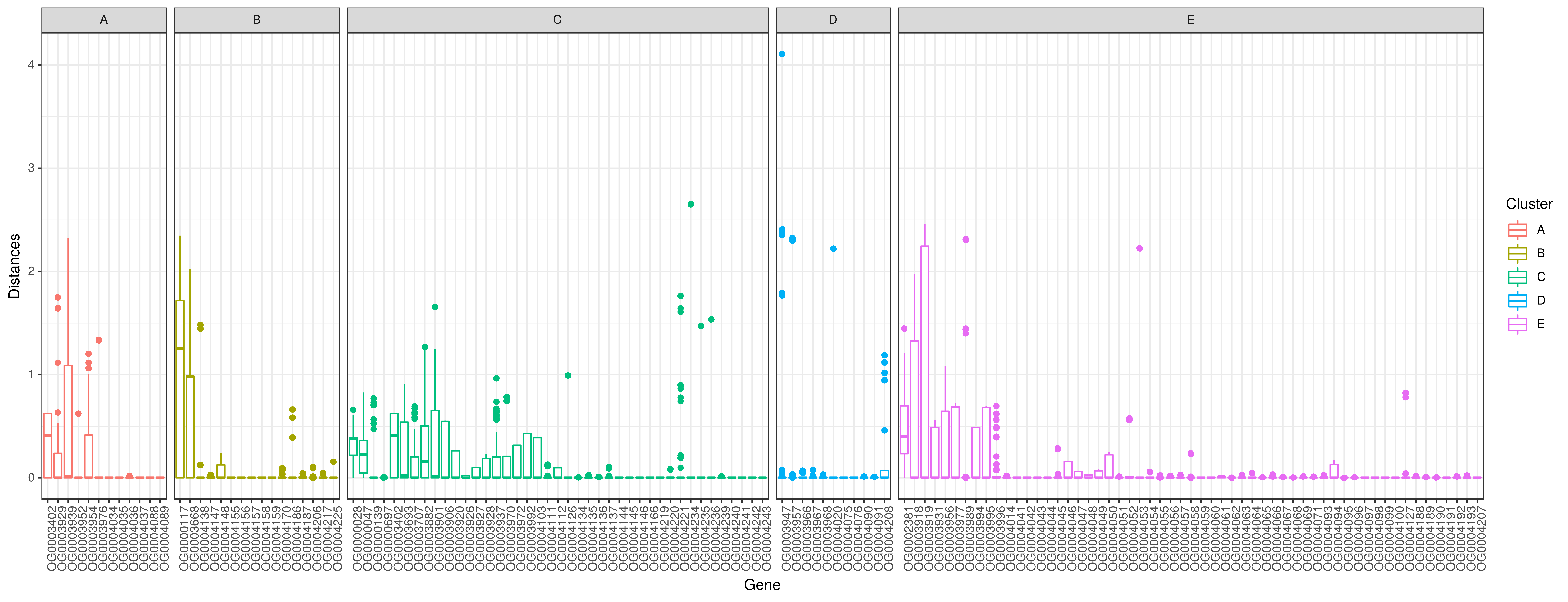

### Supplementary Figure 10

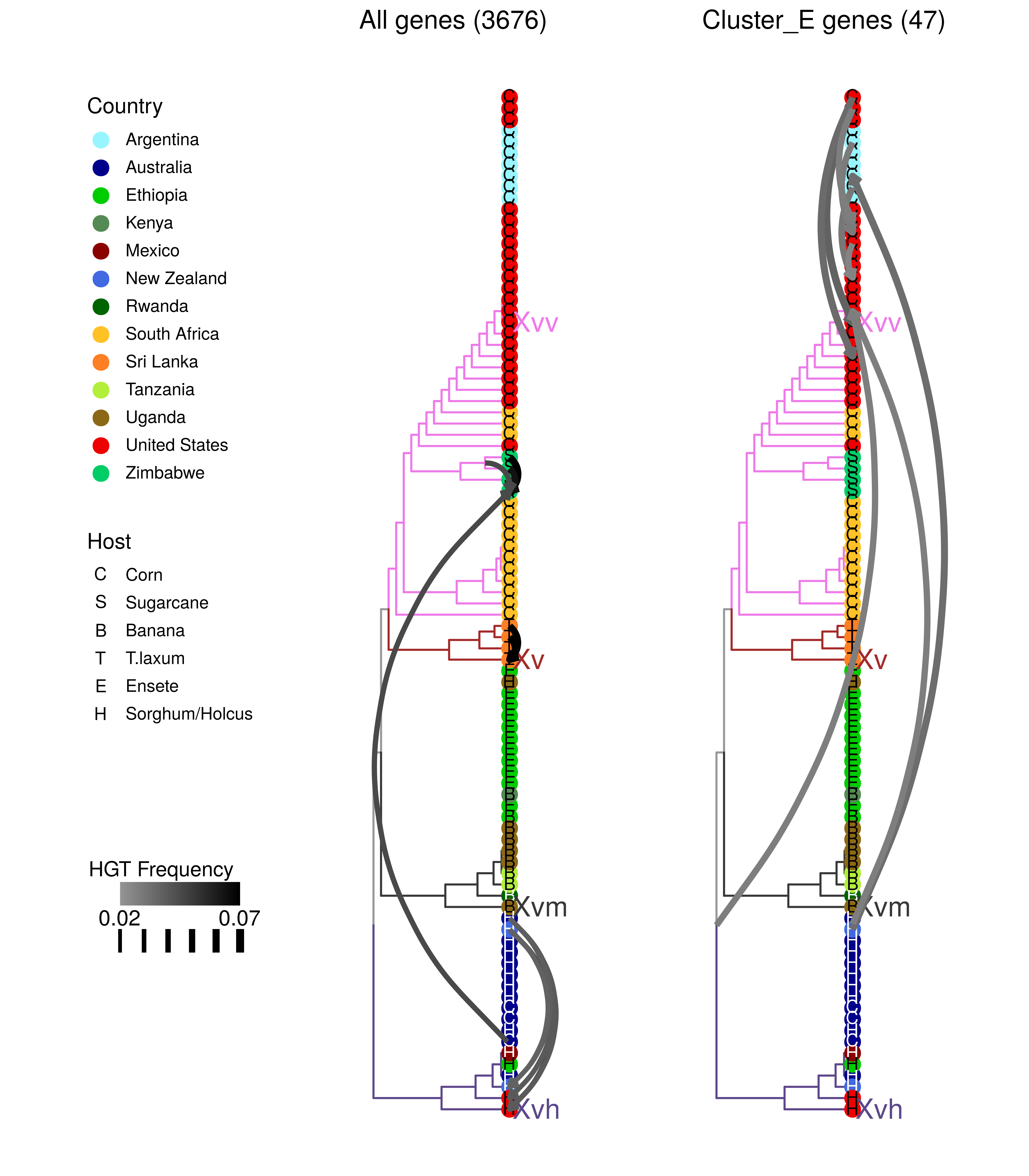

### Supplementary Figure 11

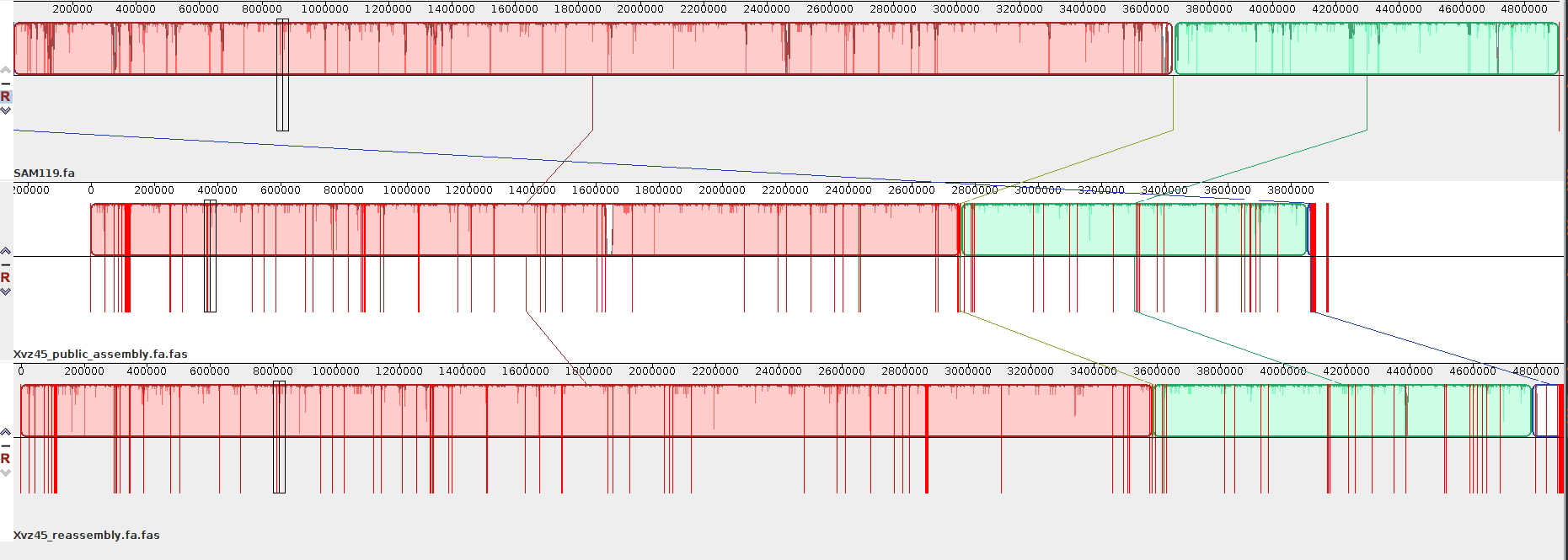
